# The first eukaryotic kinome tree illuminates the dynamic history of present-day kinases

**DOI:** 10.1101/2020.01.27.920793

**Authors:** Leny M. van Wijk, Berend Snel

## Abstract

Eukaryotic Protein Kinases (ePKs) are essential for eukaryotic cell signalling. Several phylogenetic trees of the ePK repertoire of single eukaryotes have been published, including the human kinome tree. However, a eukaryote-wide kinome tree was missing due to the large number of kinases in eukaryotes. Using a pipeline that overcomes this problem, we present here the first eukaryotic kinome tree. The tree reveals that the Last Eukaryotic Common Ancestor (LECA) possessed at least 92 ePKs, much more than previously thought. The retention of these LECA ePKs in present-day species is highly variable. Fourteen human kinases with unresolved placement in the human kinome tree were found to originate from three known ePK superfamilies. Further analysis of ePK superfamilies shows that they exhibit markedly diverse evolutionary dynamics between the LECA and present-day eukaryotes. The eukaryotic kinome tree thus unveils the evolutionary history of ePKs, but the tree also enables the transfer of functional information between related kinases.

## Introduction

Kinases are fundamental to convey information in living organisms. Due to their importance for health and agriculture, they are studied in a wide variety of eukaryotic species^1–6^. In eukaryotes, the vast majority of kinases belongs to a single family: the eukaryotic Protein Kinases (ePKs)^7, 8^. EPKs are characterised by a conserved kinase domain of about 250 amino acids that consists of 12 subdomains^9^. They phosphorylate either serine/threonine or tyrosine residues or have dual specificity. EPKs are subdivided into seven superfamilies: AGC, CAMK, CK1, CMGC, STE, Tyrosine Kinase (TK) and Tyrosine Kinase-Like (TKL)^7, 9^. EPKs that lack a superfamily have previously been referred to as ‘Other’^7^, while in this paper they will be referred to as Unaffiliated.

In 2002, a paper on the protein kinase complement of the human genome was published^7^. This highly cited paper was accompanied by an iconic poster with a phylogenetic tree of all human ePK domains. Kinome analyses of several other eukaryotic species, sometimes including a species-specific kinome tree, followed^10–14^. Currently, genomic data is available for a broad diversity of eukaryotic species. However, available kinome trees never incorporated how kinases are related across the entire eukaryotic tree of life. This is unfortunate as a eukaryotic kinome tree is relevant both to understand the function of particular ePKs better and to reveal the ancient evolutionary history of ePKs.

The functional relevance of a eukaryotic kinome tree lies foremost in facilitating the transfer of functional information between neighbouring proteins. Proteins that are close to each other in a phylogenetic tree potentially share conserved molecular interactions and mechanical properties that arose in their common ancestor. Therefore, information about well-studied ePKs in a eukaryotic kinome tree can be used to generate hypotheses for the study of related ePKs. Even ePKs that are not included in a eukaryotic kinome tree can benefit from the transfer of functional information: proteins that form a single branch in a phylogenetic tree enable classifying proteins outside the tree using Hidden Markov Models (HMMs).

A eukaryotic kinome tree is also important for the evolutionary cell biology of eukaryotes^15^. For example, current estimates of the number and identity of kinases that were already present in the Last Eukaryotic Common Ancestor (LECA) are only based on limited sets of eukaryotic species^7, 16^. A eukaryotic kinome tree allows to determine the ePK complement in the LECA more precisely.

Although a eukaryotic kinome tree is relevant both from a functional and an evolutionary perspective, there is one substantial hurdle: the large number of kinases in eukaryotes. Only the human kinome tree consists already of 491 ePK domains^7^, and in a collection of nearly 100 eukaryotes, this number increases to over 36,000 ePK domains. Such a number of sequences precludes the use of state-of-the-art alignment as well as tree building software. Moreover, it is a Sisyphean task to analyse a phylogenetic tree that consists of over 36,000 leaves.

A more general problem of gene trees is the negative impact of rapidly evolving sequences on statistic support. A commonly used strategy to improve statistic support in species trees is to select slowly evolving genes or positions^17^. For gene trees, an equivalent of this strategy has been proposed: the Scrollsaw method^18^. The Scrollsaw method systematically selects only slowly evolving sequences for generating a gene tree. As a result, both the number of sequences is reduced, and rapidly evolving sequences are excluded. This makes the Scrollsaw method perfectly suitable to handle the large number of ePKs and generate a well-supported eukaryotic kinome tree.

Here we present the first eukaryotic kinome tree, generated with a modified and extended version of the Scrollsaw method. The tree reveals ePK superfamily membership for several ePKs that have been labelled as Unaffiliated in the human kinome tree, most notably CAMKK1 and CAMKK2. The tree furthermore unveils that the LECA had much more ePKs than was thought before: at least 92. These 92 ePKs include some surprising examples of ePKs that were previously believed to be specific for certain eukaryotic supergroups, like human CHK1 and the plant CIPKs. The number of LECA ePKs retained in present-day species varies enormously within and between eukaryotic supergroups. The expansion of LECA ePKs since the common ancestor of eukaryotes also differs within and between ePK superfamilies. This variation in LECA ePK dispensability and duplicability is possibly linked to differential roles of LECA ePKs in cellular housekeeping and organismal innovation. The eukaryotic kinome tree thus both reveals the evolutionary history of ePKs and directs the study of ePK function.

## Results

### The eukaryotic kinome tree reveals well-supported LECA kinase clades

In order to generate a eukaryotic kinome tree, 36,475 ePK domains were collected from 94 eukaryotic species (Supplementary Data 1). These domains were used as input for a pipeline that generates two phylogenetic trees by implementing a modified and extended version of the Scrollsaw method (Fig. 1, Methods). In this LECA clade annotation pipeline, the Scrollsaw method was extended with automatic annotation of the tree leaves into LECA kinase clades: groups of kinases that likely have a single ancestor in the LECA because they include at least one Amorphea and one Bikonta sequence^19^. The LECA clade annotation pipeline generated two different eukaryotic kinome trees in order to facilitate automatic annotation, cross-validate annotated LECA kinase clades and produce HMM profiles that contain more sequence diversity. One tree is based on Bi-directional Best Hits (BBHs) between two eukaryotic supergroups (Fig. 2, Supplementary Data 2-5), while the other tree is based on BBHs between five eukaryotic supergroups (Supplementary Fig. 1, Supplementary Data 6-9). The leaves of both trees were automatically annotated into LECA kinase clades, and both sets of LECA kinase clades were combined into one non-overlapping set. This combined set was improved manually, resulting in a final set of 118 LECA kinase clades.

**Fig. 1:**
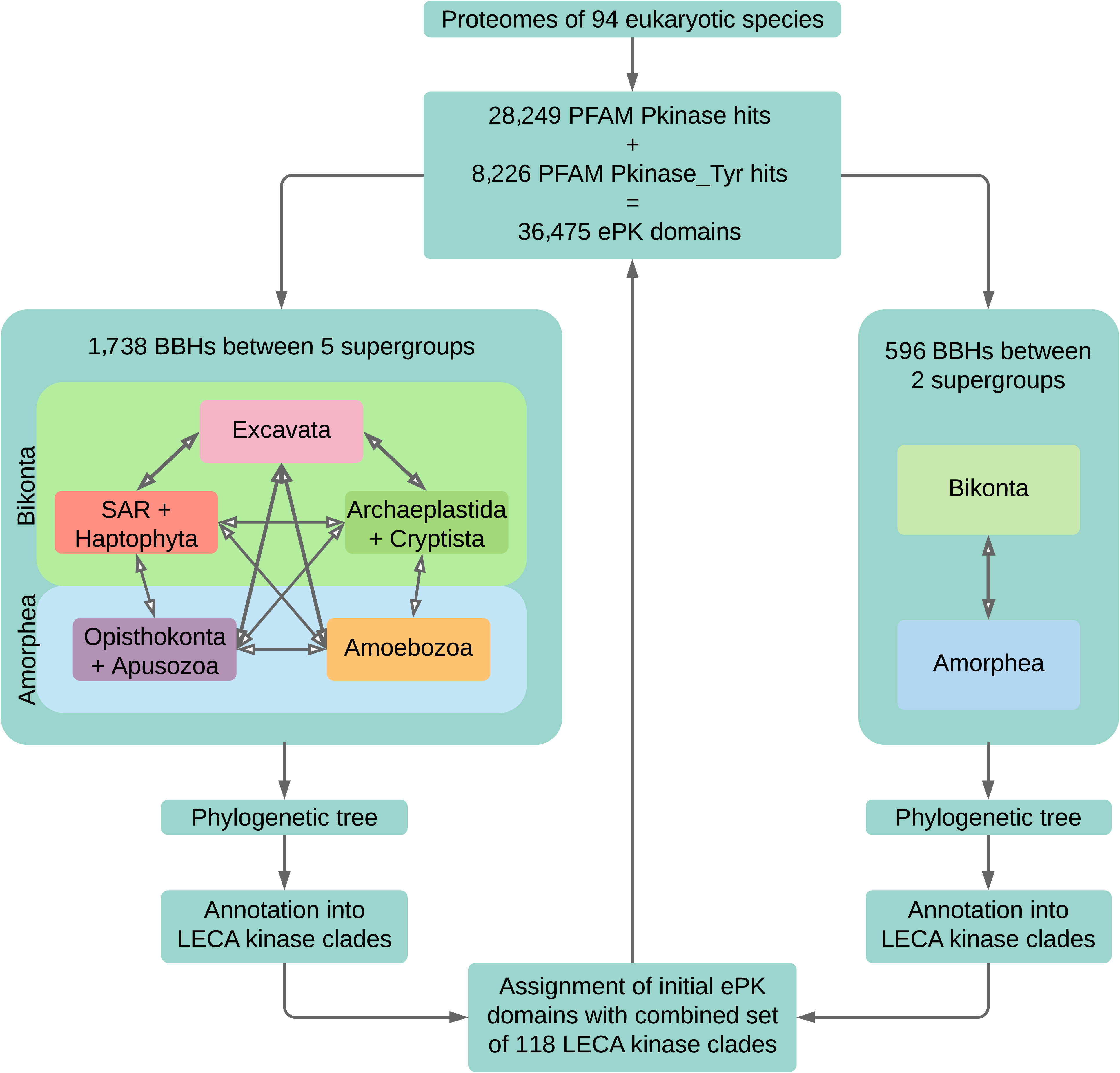
Overview of important steps in the LECA clade annotation pipeline.

**Fig. 2:**
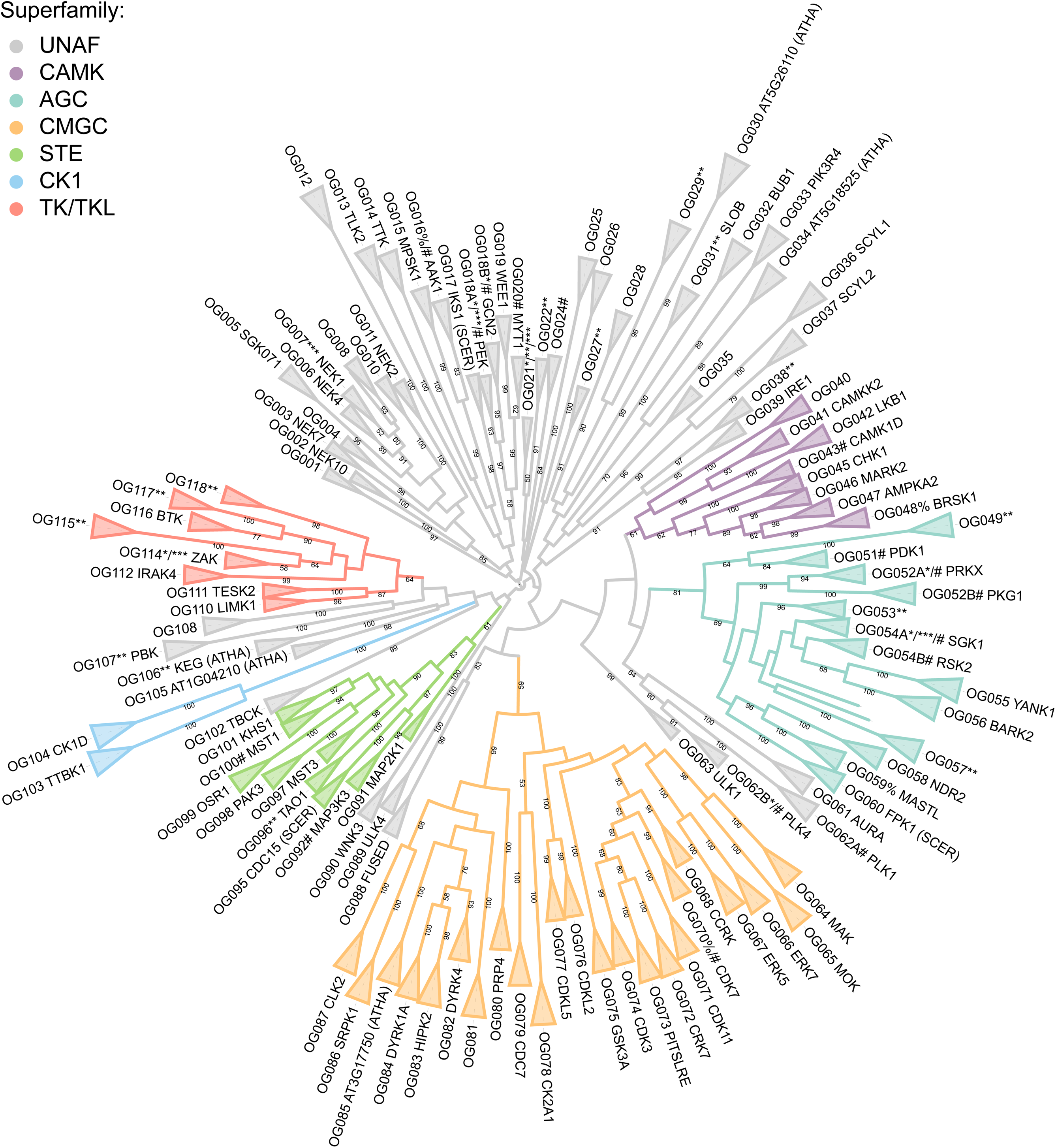
The eukaryotic kinome tree based on two-supergroups-BBHs. LECA kinase clades are colour coded according to ePK superfamily. LECA kinase clade names indicated with SCER or ATHA in-between brackets are not derived from human but from baker’s yeast or *A. thaliana* protein names, respectively. Unaffiliated LECA kinase clades are grey. LECA kinase clades that fail one or more criteria for inclusion in the LECA kinase number estimate are indicated with *(absence in one of the trees), **(limited distribution over species) and ***(bootstrap support below 70 in both trees). LECA kinase clades that initially were split into two LECA kinase clades in one of the trees are indicated with %. LECA kinase clades that include manually annotated leaves are indicated with #. Bootstrap support of minimal 50 out of 100 is shown.

Even though the LECA lived about 1-1.9 billion years ago^20^, the vast majority of LECA kinase clades in the eukaryotic kinome tree is statistically well supported. For example, in the two-supergroups-BBHs tree, 78 per cent of the LECA kinase clades have bootstrap support values of 95 or above (Fig. 2). This percentage is lower for the five-supergroups-BBHs tree (Supplementary Fig. 1). Higher support for LECA kinase clades in the two-supergroups-BBHs tree is in agreement with results from the original Scrollsaw paper^18^. There, bootstrap support for LECA clades increased upon additional reduction of BBHs to one sequence per eukaryotic supergroup per LECA clade. The high bootstrap support for our LECA kinase clades confirmed the usefulness of the Scrollsaw method to obtain well-supported orthologous clades in large gene families.

Although LECA kinase clades are well supported, internal support in the eukaryotic kinome tree is lower (Supplementary Results). Bootstrap support values of at least 70 are found only in 26 per cent of the 112 pre-LECA duplications in the two-supergroups-BBHs tree (Fig. 2). Surprisingly, 34 per cent of these well-supported pre-LECA kinase clades appear deeper in the tree and include three or more LECA kinase clades.

### Nearly 80 per cent of kinase domains can be assigned to a LECA kinase clade

Protein assignment via HMM profiles outperforms assignment via BLAST^21^. Therefore, HMM profiles of the 118 LECA kinase clades of the eukaryotic kinome tree (Supplementary Data 10) were used to automatically assign the initial set of 36,475 kinase domains (Fig. 1, Supplementary Table 1, Supplementary Data 11). Despite a conservative approach, 28,893 kinase domains were assigned to their top hitting LECA kinase clade. Kinase domains were not assigned if the top two scoring LECA kinase clades had a bit score difference below 10 or a maximal bit score below 30 (Supplementary Data 12 and 13). Although the assigned kinase domains encompass nearly 80 per cent of the total kinase domain dataset, there is considerable variation in assignment percentages between species (Supplementary Table 2). For each LECA kinase clade, the protein name of the best scoring human kinase domain was used to name the LECA kinase clade. If human hits were not available, best hits from baker’s yeast or *Arabidopsis thaliana* were used for naming. LECA kinase clades to which no kinase domains from these three species were assigned are indicated with Orthologous Group (OG) and a number.

### LECA complexity involved at least 92 eukaryotic protein kinases

Our reconstruction of LECA kinase clades enabled estimating the LECA kinase repertoire. However, not all 118 LECA kinase clades are equally likely to represent a single gene in the ancestor of all eukaryotes. An initial conservative estimate of the most reliable LECA kinase clades yielded 91 ePKs in the LECA. These 91 LECA kinases correspond to 91 LECA kinase clades that are annotated in both eukaryotic kinome trees (Fig. 2, Supplementary Fig. 1), have at least one bootstrap support of minimal 70, and kinase domains from minimal two eukaryotic supergroups have been assigned to them (Supplementary Data 14). Other LECA kinase clades did not fulfil one or more of these criteria and require more investigation. For example, LECA kinase clades to which only a limited number of kinase domains were assigned need closer examination. This could discriminate between possible explanations like horizontal gene transfer, genome contamination, or being a bonafide LECA kinase that has been lost in many species.

In addition to the 91 LECA ePKs that are based on LECA kinase clades, one more LECA kinase was added: Haspin. This kinase was absent in our kinase domain dataset because it is a diverged ePK with a PFAM model distinct from the PFAM models that were used to collect sequences for the eukaryotic kinome tree^8^. Haspin was likely already present in the LECA^16, 22^. Thus our initial conservative estimate of 91 ePKs in the LECA together with Haspin result in an estimate of 92 ePKs in the LECA. This is more than a third more than the largest previous estimate of 68 basal ePKs^16^ (Supplementary Results). A large LECA ePK complement is in line with a LECA that was much more complex than many present-day eukaryotes^20^.

### The common ancestry of LKB1 and CAMKKs explains their functional overlap

The eukaryotic kinome tree is highly consistent with the human kinome tree, but a complete agreement would require some adjustments to the human kinome tree (Supplementary Results). The eukaryotic kinome tree, for example, clarified the relationships between a few Unaffiliated human kinases and the ePK superfamilies (Supplementary Results). The most interesting example of Unaffiliated human kinases that stem from within an ePK superfamily are the human kinases assigned to LECA kinase clade CAMKK2 (Fig. 2, Supplementary Fig. 1). The names of these kinases, CAMKK1 and CAMKK2, already reflect their functional link with the CAMK superfamily. Despite this functional link, CAMKK1 and CAMKK2 were placed at a distance from the CAMK superfamily in the human kinome tree (Supplementary Fig. 2). In contrast, LECA kinase clade CAMKK2 is positioned within the CAMK superfamily in the eukaryotic kinome tree. It is located next to LECA kinase clade LKB1 with high bootstrap support (99; Fig. 2).

The juxtaposition of LECA kinase clades LKB1 and CAMKK2 is striking both from a functional and evolutionary perspective. In human, LKB1 is a master kinase of AMPK and the AMPK-related kinases^23^, which were assigned to the LECA kinase clades AMPKA2, MARK2 and BRSK1 (Supplementary Table 1). In addition to LKB1, AMPK can also be phosphorylated by CAMKK2^24^. In baker’s yeast and *A. thaliana*, orthologs of CAMKK1 and CAMKK2 are the only upstream kinases of AMPK orthologs because LKB1 is absent in these species^25^ (Supplementary Table 1). The juxtaposition of LECA kinase clades LKB1 and CAMKK2 is therefore in agreement with their overlapping function in AMPK phosphorylation. It suggests that the common ancestor of LECA kinase clades LKB1 and CAMKK2 may already have been able to phosphorylate the common ancestor of LECA kinase clades AMPKA2, MARK2 and BRSK1. The common ancestor of LKB1, CAMKK2 and possibly OG040 may even have been able to phosphorylate the common ancestor of all other CAMK LECA kinase clades. This is suggested by the basal position of LKB1, CAMKK2 and OG040 within the CAMK superfamily (Fig. 2, Supplementary Fig. 1), and the fact that CAMKK2 can also phosphorylate members of LECA kinase clade CAMK1D^26^. Interestingly, also within the AGC superfamily, the most basal position is reserved for a master kinase: PDK1^27^ (Fig. 2, Supplementary Fig. 1).

### The majority of Unaffiliated human kinases stem from Unaffiliated kinase clades

One of our reasons to generate the eukaryotic kinome tree was to test whether there exist Unaffiliated human kinases that actually belong to an ePK superfamily. Including CAMKK1 and CAMKK2, 14 of the 88 Unaffiliated human kinases could be classified into a superfamily (Supplementary Results). However, no less than 56 Unaffiliated human kinases were assigned to 17 Unaffiliated (pre-)LECA kinase clades that have no well-supported further affiliations in the eukaryotic kinome tree (Supplementary Results). Although their phylogenetic position might be insufficiently resolved due to accelerated evolution, many of the Unaffiliated (pre-)LECA kinase clades could also be the result of old duplications early in eukaryogenesis. Such an old age could explain why it is difficult to connect Unaffiliated (pre-)LECA kinase clades firmly to any other (pre-)LECA kinase clade or ePK superfamily. The Unaffiliated (pre-)LECA kinase clades are then as old and distinct as entire ePK superfamilies but duplicated less vigorously during eukaryogenesis than most ePK superfamilies.

### Human CHK1 and plant CIPKs were one kinase in the LECA

The LECA kinase clade delineation suggested a single LECA ancestor for kinases from different eukaryotic groups that so far were thought to be group-specific. A prominent case is the common ancestry of plant CBL-Interacting Protein Kinases (CIPKs) and opisthokont Checkpoint Kinase 1 (CHK1). CIPKs, also known as SNRK3s, have often been described as plant-specific, but recently they have also been found in other eukaryotic species^28^. CHK1 is considered opisthokont-specific^29^. However, both kinase families were assigned to LECA kinase clade CHK1 within the CAMK superfamily, suggesting a common ancestor in the LECA (Supplementary Table 1, Fig. 2, Supplementary Fig. 1). This suggestion is supported by some additional experimental and phylogenetic evidence^21, 29–33^.

The CIPK-specific NAF domain is absent in opisthokont CHK1. However, this domain is still present in *Thecamonas trahens* CHK1 (Supplementary Data 15). *T. trahens* is a member of the Apusozoa, the sister clade of opisthokonts, and therefore belongs to the Amorphea. The presence of the NAF domain in both Amorphea CHK1 and Bikonta CIPKs suggests that the LECA CHK1 had a NAF domain as well. The absence of the NAF domain in opisthokont CHK1 is, therefore, a derived feature due to a loss in the common ancestor of fungi and animals.

### Human HIPKs and baker’s yeast YAK1 were one kinase in the LECA

Another unrecognised deep orthology was found between the human Homeodomain-Interacting Protein Kinases (HIPKs) and baker’s yeast Yet Another Kinase 1 (YAK1). YAK1 orthologs are present throughout eukaryotes, but they were thought to be missing in Metazoa^16^. In KinBase^34^, the HIPK and YAK subfamilies have a perfectly complementary distribution: either Metazoa-specific or missing in Metazoa. However, both human HIPKs and baker’s yeast YAK1 were assigned to LECA kinase clade HIPK2 within the CMGC superfamily (Supplementary Table 1, Fig. 2, Supplementary Fig. 1).

In earlier studies, the HIPKs and YAK1 have been suggested to be different classes of DYRK proteins^35, 36^. However, depending on how the phylogenetic trees in those studies are rooted, the HIPKs and YAK1 could be inferred to be monophyletic. HIPK2 and YAK1 also have a shared function in phosphorylating the CCR4-NOT complex^37^. Together these data suggest that the metazoan HIPKs and the eukaryote-wide found YAK1 were indeed one kinase in the LECA. The fact that metazoan HIPKs apparently are difficult to recognise as YAK1 orthologs indicates derived characteristics for HIPKs.

### The LECA kinase presence-absence profile displays diverse patterns of kinase retention

The assignment of the initial set of 36,475 ePK domains to LECA kinase clades allowed the generation of a clustered presence/absence profile of LECA kinases in present-day species (Fig. 3). This clustering divided LECA kinases into two large clusters. The ‘ubiquitous’ cluster at the top of the presence/absence profile contains 49 LECA kinases, of which 48 are present in at least half of the 94 species. Four LECA kinases are retained in all extant eukaryotic species in our dataset: CDK3 (but note that CDK3 might comprise two nested LECA kinase clades, see Supplementary Results), CAMK1D, CK2A1 and CK1D. Several other LECA kinases are nearly omnipresent: AURA, CRK7, GSK3A, ERK5, MAP2K1, CAMKK2, SRPK1, PDK1, NDR2 and AMPKA2.

**Fig. 3:**
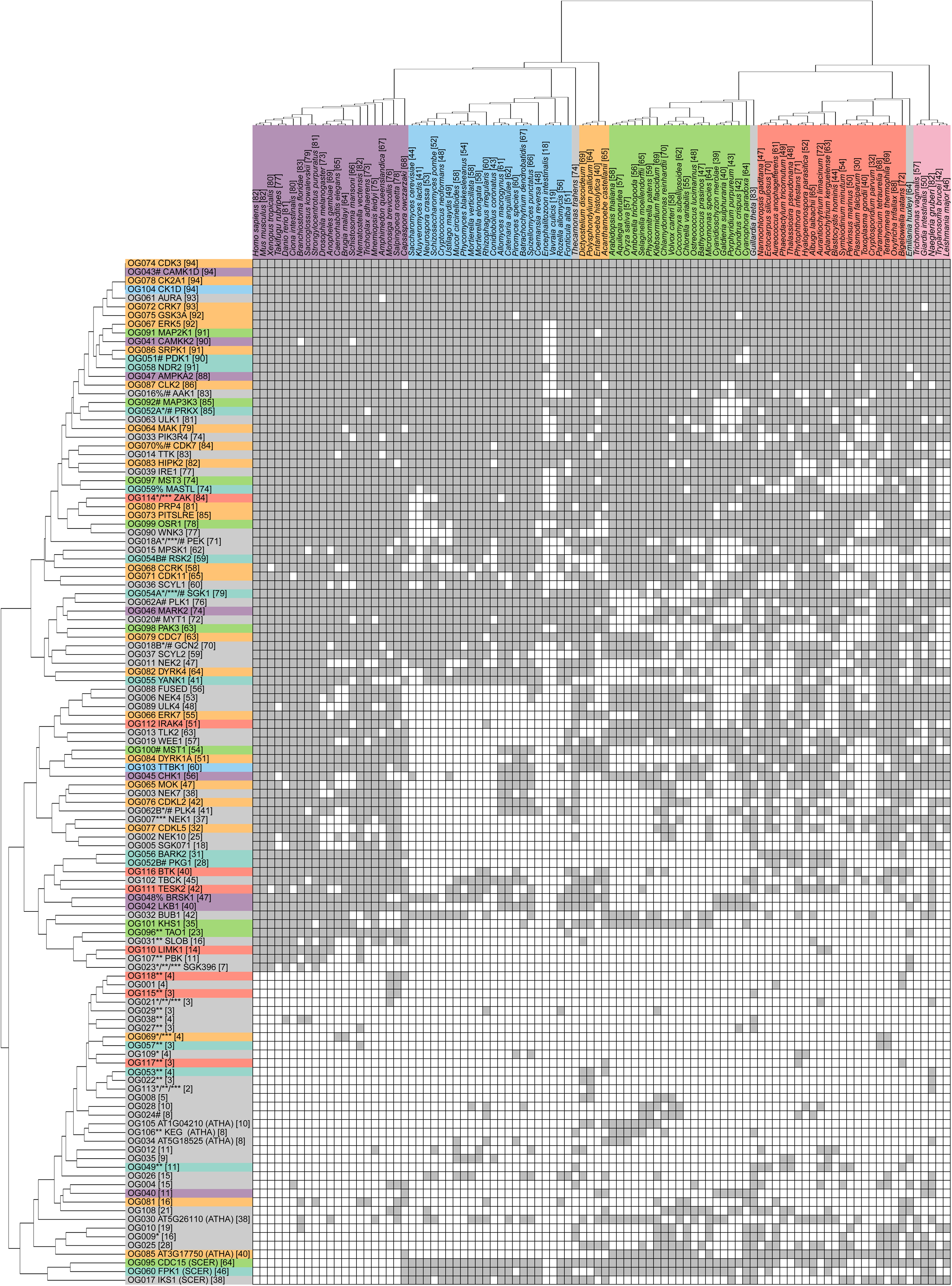
Clustered presence/absence profile of 28,893 ePK domains in 94 present-day eukaryotes. Assignment of one or more ePKs from a species to a certain LECA kinase clade is indicated in grey while absence is indicated in white. LECA kinase colour code and the meaning of special characters is the same as in Fig. 2. Species are colour coded according to eukaryotic supergroup as in Fig. 4. Behind LECA kinases the total number of species from which at least one ePK was assigned to a particular LECA kinase clade is given. Behind species names the total number of LECA kinase clades to which ePKs from a particular species were assigned is given.

The cluster at the bottom of the presence/absence profile can be subdivided into two subclusters. The first ‘fungal-loss’ subcluster contains 33 LECA kinases, of which 13 are present in at least half of the 94 species, just like the kinases in the ‘ubiquitous’ cluster. However, many of the LECA kinases in the ‘fungal-loss’ subcluster were lost in several or all fungi. The second ‘sparse’ subcluster contains 36 LECA kinases, of which 30 are present in less than a quarter of the 94 species. Not surprisingly, the ‘sparse’ cluster encompasses the majority of LECA kinases that were excluded from the LECA kinase number estimate because they were not present in a sufficient number of species (indicated with **).

Within the ‘sparse’ subcluster, OG040 is particularly interesting. This LECA kinase is nearly only found in early-branching species: two excavates, *Guillardia theta*, *Cyanophora paradoxa,* red algae, two amoebae and *Capsaspora owczarzaki*. OG040 also has an interesting position in the eukaryotic kinome tree: it clusters next to the master kinase clade that contains CAMKK2 and LKB1 (Fig. 2, Supplementary Fig. 1). The retention pattern and phylogenetic position of OG040 raise curiosity about its function in the LECA and in present-day species.

### LECA kinase retention varies between and within eukaryotic supergroups

The differential presence of eukaryotic supergroups in the ‘ubiquitous’, ‘fungal-loss’ and ‘sparse’ cluster is reflected in Fig. 4, where species are ordered by their total number of LECA kinases. Holozoa, a group that comprises animals and their unicellular relatives, dominate the top of this graph. They are headed by *Branchiostoma floridae*, which shares the maximum number of 83 retained LECA kinases with the cryptophyte *G. theta*. Early-branching unicellular Holozoa are not found among the top-scoring Holozoa, but the Choanomonadida *Salpingoeca rosetta* and *Monosiga brevicollis* still kept 78 and 76 LECA kinases respectively. *C. owczarzaki* kept only 68 LECA kinases and lost quite some LECA kinases that are present in most Holozoa. However, it also retained several LECA kinases that were lost in other Holozoa, like CDC15, FPK1 and IKS1.

**Fig. 4:**
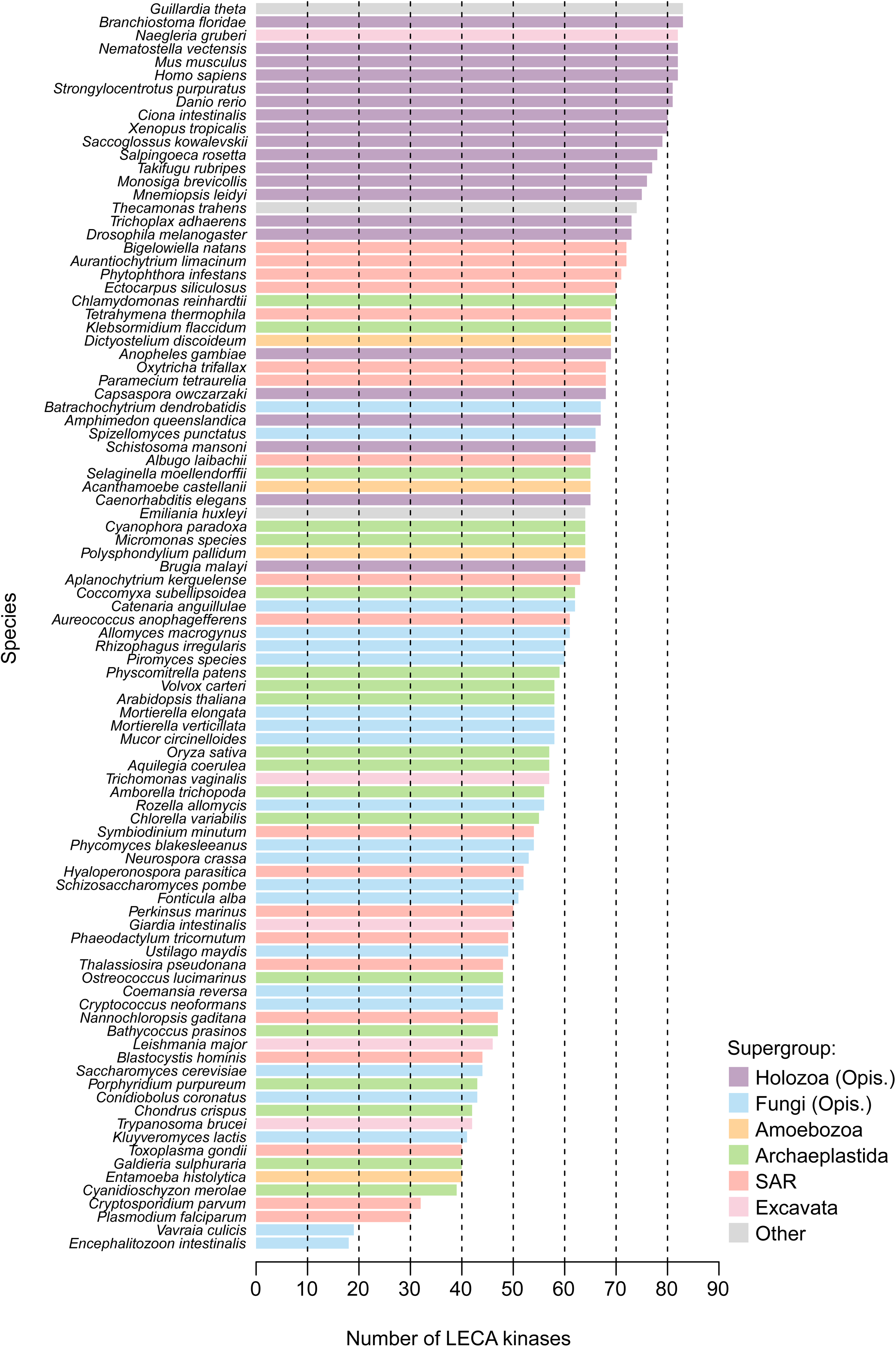
LECA kinase retention in 94 present-day eukaryotes. Species are colour coded according to eukaryotic supergroup.

The other large group within the opisthokonts, the fungi, exhibit a pattern strikingly different from the Holozoa: they are mainly present at the bottom of Fig. 4. The fungi that kept most LECA kinases are relatively early-branching species like the Chytridiomycota *Batrachochytrium dendrobatidis* (67 LECA kinases) and *Spizellomyces punctatus* (66 LECA kinases). Within the Archaeplastida, also early-branching species like the green alga *Chlamydomonas reinhardtii* and charophyte alga *Klebsormidium flaccidum* kept the largest number of about 70 LECA kinases. Interestingly, the model organisms *Saccharomyces cerevisiae* and *A. thaliana* maintained a relatively small number of respectively 44 and 58 LECA kinases. At the very bottom of Fig. 4, intracellular parasites with streamlined genomes like the fungal Microsporidia *Vavraia culicis* and *Encephalitozoon intestinalis* are found^38^. They kept less than 20 LECA kinases.

The small numbers of 39-43 LECA kinases that have been retained in the red algae *Chondrus crispus*, *Galdieria sulphuraria*, *Cyanidioschyzon merolae* and *Porphyridium purpureum* can probably also be attributed to a genome reduction^39^. However, the red algae, just like *C. owczarzaki* within the Holozoa, also reflect their early-branching position within the Archaeplastida. Together with *C. paradoxa,* they kept three LECA kinases that were lost in the rest of the Archaeplastida lineage. These three LECA kinases, LKB1, its neighbour OG040 and its downstream kinase BRSK1, are all members of the CAMK superfamily. A fourth LECA kinase, PAK3 from the STE superfamily, is kept only in red algae but not in *C. paradoxa*. Interestingly, a human kinase assigned to LECA kinase clade PAK3 is inhibited by LKB1^40^. Therefore all four LECA kinases that have been retained only in basal Archaeplastida are phylogenetically or functionally related. This suggests that in these species, they may participate in the same process.

### The largest LECA kinase superfamily CMGC expanded least from LECA to human

The eukaryotic kinome tree together with the presence-absence profile of its LECA kinases enabled to quantify the evolutionary dynamics of ePK superfamilies from LECA till present-day species. Except for the CMGC and CK1 superfamilies, ePK superfamilies had about 10 members in the LECA (Fig. 2, Supplementary Fig. 1, Fig. 5). The CMGC superfamily was much larger with 24 LECA kinases while the CK1 superfamily was much smaller with two kinases in the LECA. Although most kinase superfamilies had comparable sizes in the LECA, their expansion from LECA to human is strikingly different (Fig. 5). The large CMGC superfamily expanded 2.8 times from LECA to human. The small CK1 superfamily and medium-sized STE and AGC superfamilies are about five times larger in human compared to the LECA. The other medium-sized superfamilies expanded about 10 times (CAMK) or even more (TK/TKL).

**Fig. 5:**
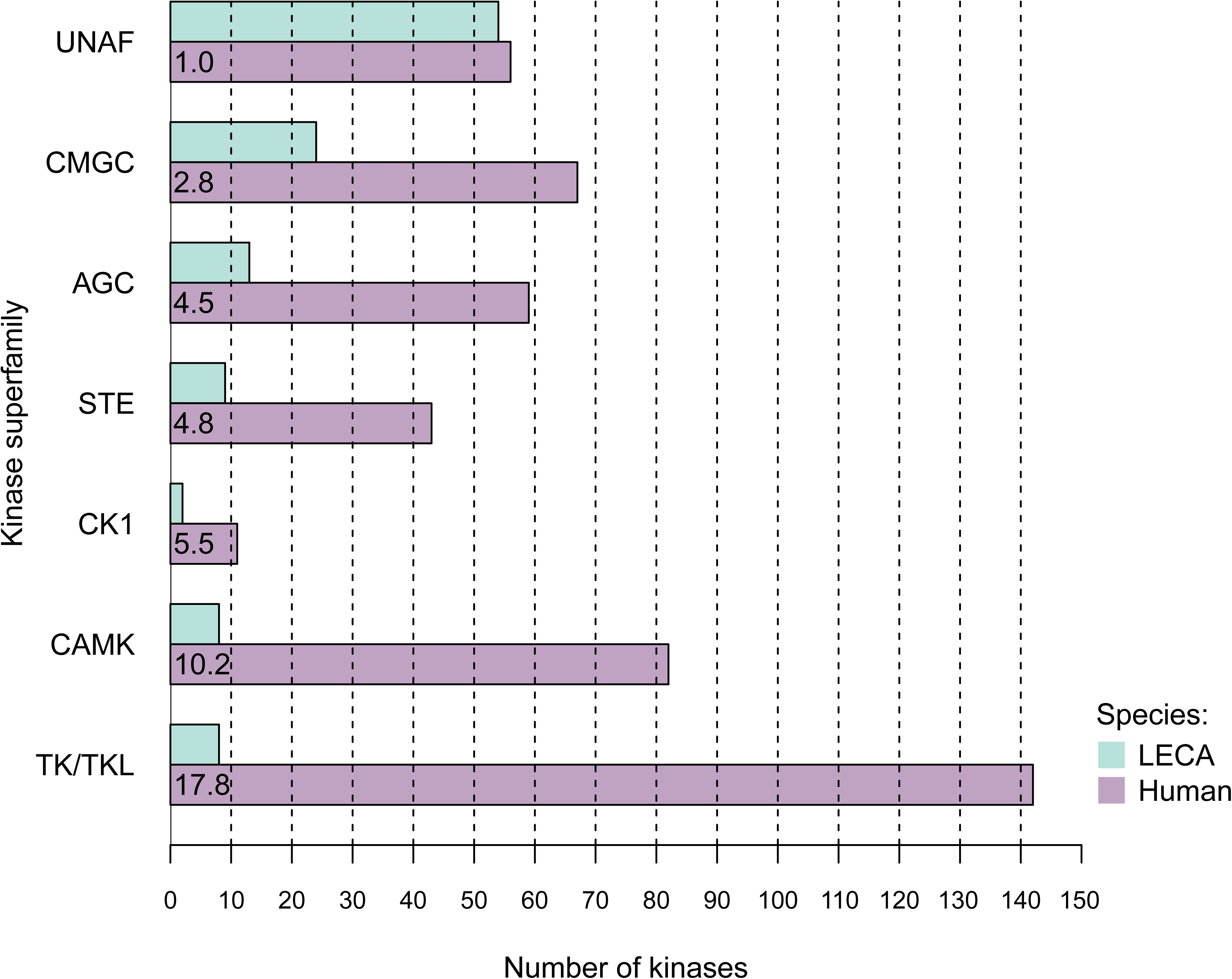
The expansion of ePK superfamilies from LECA till human. Human bars show the multiplication factor between the number of kinases in the LECA and in human.

### Kinase superfamily duplicability and dispensability are variable

In general, superfamily expansion from LECA to human and the average expansion of single LECA kinases from that same superfamily in 94 eukaryotes display a similar trend (Fig. 5, Fig. 6). For example, LECA kinases from the large CMGC superfamily did not expand much from LECA to human, and their per LECA kinase average expansion in 94 eukaryotes is also low. However, kinase expansion from LECA to human and other present-day species is also variable within superfamilies.

**Fig. 6:**
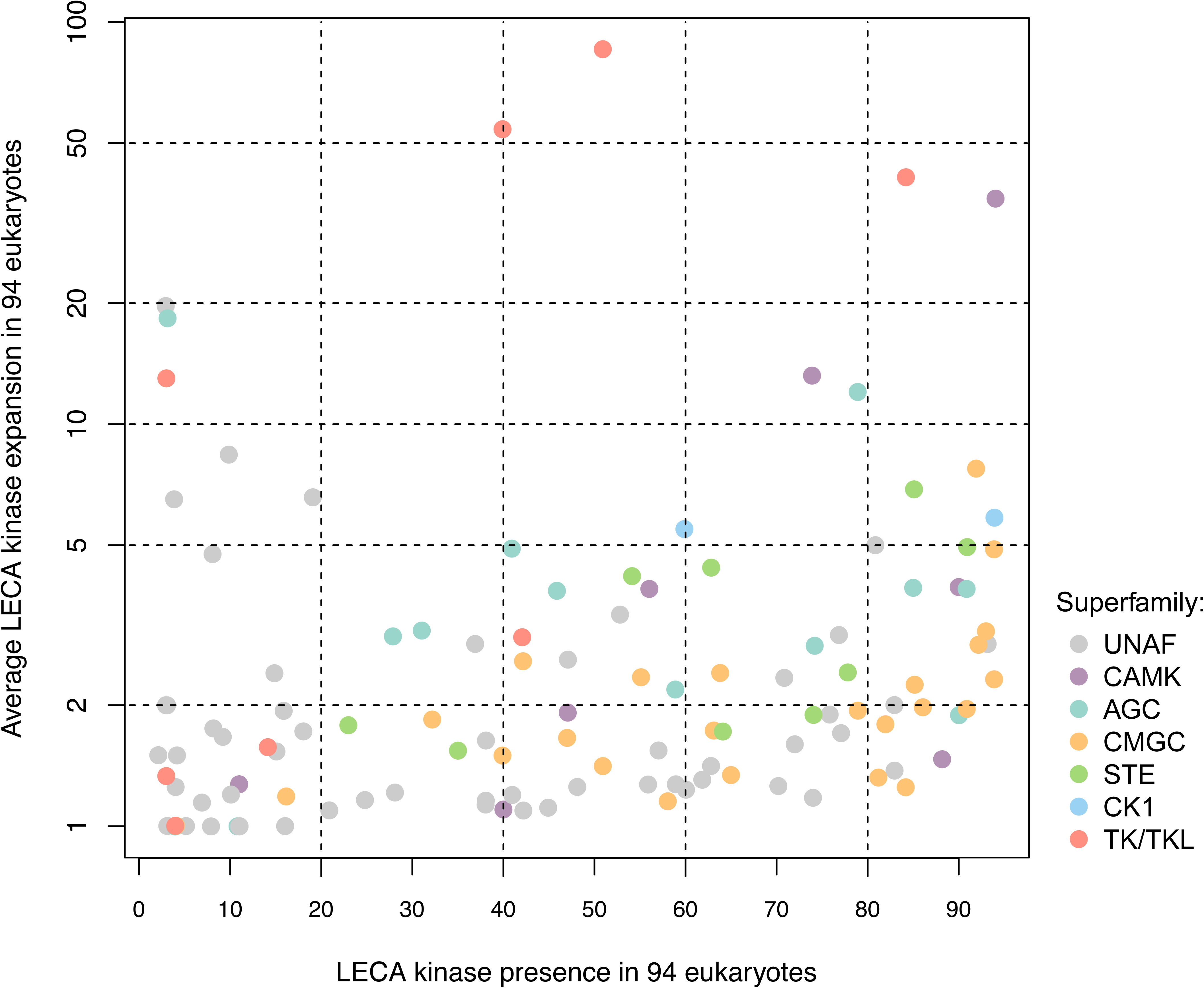
EPK superfamily dynamics. The presence of 118 LECA kinases in 94 present-day eukaryotes is shown versus the average expansion of these LECA kinases in the same set of species. LECA kinases are colour coded according to ePK superfamily.

The most striking variation in LECA kinase expansion is found within the TK/TKL and CAMK superfamilies. These superfamilies expanded most from LECA to human (Fig. 5). However, the average gene count of TK/TKL and CAMK LECA kinases in present-day eukaryotes is predominantly low (<2) or high (>10) (Fig. 6). Some TK/TKL and CAMK LECA kinases, including TK/TKL LIMK1 and CAMK AMPKA2, hardly expanded from LECA to present-day eukaryotes. These kinases are likely to perform similar functions in extant eukaryotes as in the LECA. On the other hand, LECA kinases like TK/TKL IRAK 4 and CAMK CAMK1D, underwent much duplication. Descendants of these LECA kinases likely perform various new functions.

LECA kinases from the CMGC superfamily are special: they exhibit together with low duplicability also low dispensability (Fig. 6). The average expansion of CMGC kinases from LECA to present-day eukaryotes is just above 2, while nearly half of the CMGC LECA kinases are present in more than 80 present-day eukaryotes. LECA kinases from the TK/TKL superfamily display in contrast to the CMGC superfamily both high duplicability and high dispensability (Fig. 6).

### Unaffiliated LECA kinases display low duplicability

In the eukaryotic kinome tree, a large group of 53 LECA kinases is not part of any of the superfamilies (Fig. 2, Supplementary Fig. 1, Fig. 5). A comparison with the human kinome tree suggested that most of these Unaffiliated LECA kinases duplicated infrequently. The expansion of the Unaffiliated LECA kinases from LECA to human is indeed very low, even below that of the CMGC kinases (Fig. 5). However, the Unaffiliated LECA kinases and the CMGC superfamily display different relationships between duplicability and dispensability. The Unaffiliated LECA kinases generally combine low duplicability like the CMGC LECA kinases with high dispensability like the TK/TKL LECA kinases (Fig. 6). Because the Unaffiliated LECA kinases do not form a monophyletic group in the eukaryotic kinome tree, they should nevertheless not be treated as a set of kinases with similar evolutionary and functional properties. For example, several Unaffiliated LECA kinases including AAK1 and TTK duplicated infrequently but have dispensability levels comparable to LECA kinases from the CMGC superfamily.

## Discussion

We demonstrate for the first time that by using only slowly evolving kinases, it is possible to generate a eukaryotic kinome tree. The eukaryotic kinome tree was largely automatically annotated into well-supported LECA kinase clades for estimating the ePK repertoire of the LECA. Subsequently, HMM profiles of LECA kinase clades were used to assign the majority of ePKs from 94 eukaryotic genomes to the LECA kinase clade from which they most likely originated.

The eukaryotic kinome tree reveals the phylogenetic relationships between ePKs that were already present in the LECA. The tree can also be used as a platform to better understand ePK evolution on a more functional level. For example, mapping ePK functions on the eukaryotic kinome tree can be used to estimate the relative age of cellular processes that played a role in eukaryogenesis. This requires sufficient internal support to establish the duplication order of LECA kinase clades. However, we observed a puzzling contrast in support between pre-LECA kinase clades (low support) and LECA kinase clades (high support). Interestingly, the limited number of pre-LECA kinase clades that *are* well-supported often include more than two LECA kinase clades.

Pre-LECA support might improve upon further reducing the BBH set, adding non-kinase domains and advancing phylogenetic methods^41^. It is also possible that the exact order of many pre-LECA duplications will remain unsolvable. The existence of well-supported pre-LECA kinase clades that contain multiple LECA kinase clades suggests that these pre-LECA kinase clades may have undergone several rounds of rapid duplication during eukaryogenesis. Rapid duplication complicates reconstructing the duplication order within a pre-LECA kinase clade. An alternative explanation for weak internal support in pre-LECA kinase clades is a syncytial LECA^42^. In that case, presumed pre-LECA duplications are a form of alloploidy.

Based on the eukaryotic kinome tree, we estimated that the number of LECA ePKs is much larger than previously thought. At the same time, our estimate is conservative as only consistent LECA kinase clades with sufficient support values are included. Additional LECA kinase clades might be found upon the sampling of novel eukaryotic species. Especially non-photosynthetic free-living protists are under-sampled in currently sequenced eukaryotes^43^. The LECA ePK estimate might further increase upon iteratively updating the PFAM HMM profiles Pkinase and Pkinase_Tyr. Searches with the current HMM profiles perhaps resulted in some false negatives^44^ that by inclusion in the BBH sets could result in new LECA kinase clades. Finally, the number of LECA kinase clades could be expanded by manual inspection of potential LECA kinase clades that are not yet included in the current LECA ePK estimate. These potential LECA kinase clades illustrate that our automatic approach is useful at prioritising regions of the eukaryotic kinome tree where manual research is most needed.

By classifying LECA ePKs in ePK superfamilies, we revealed variation in duplicability and dispensability between and within ePK superfamilies. Orthologous genes that have a wide phyletic distribution are often essential genes^45, 46^. Therefore LECA kinases that display low dispensability may perform functions that have remained essential throughout eukaryotic evolution. LECA kinases that hardly duplicated may also have largely retained their original function from LECA till present-day eukaryotes. Although gene retention or loss after duplication is not necessarily adaptive^47^, highly duplicated genes have been connected to phenotypical changes^21^. Highly duplicated ePKs may thus have contributed to an increased regulatory potential on the evolutionary trajectory from early to present-day eukaryotes. For example, the enormous expansion of kinases from the TK/TKL superfamily in human and other metazoans was indeed essential for the development of multicellularity^48^.

The CMGC superfamily is unique, as it combines being the largest LECA kinase superfamily with low duplicability post-LECA and low dispensability in present-day species. The current low duplicability and low dispensability are likely due to essential functions that the CMGC kinases performed in the LECA and still perform in present-day species. The large number of duplications in the CMGC superfamily during eukaryogenesis, in contrast, may have allowed adaptive evolution towards the complex eukaryotic cell.

The eukaryotic kinome tree reveals various new insights in the evolution of kinases. However, the results that we present in this paper also demonstrate that the Scrollsaw method is a valuable approach to generate a well-supported phylogenetic tree starting from a large set of proteins. Our extension of the Scrollsaw method with automatic LECA clade annotation helps to analyse such a tree in a quick and reproducible way. The LECA clade annotation pipeline that we built can, in principle, be applied to all eukaryotic protein domains with sufficient length to generate phylogenetic trees.

The data that we generated are furthermore a rich resource for a more functional approach to kinases. The eukaryotic kinome tree can serve to select closest neighbours for information transfer, and the overview of LECA kinase retention is useful to select species that are most fit to study the LECA kinase repertoire. The HMM profiles that we provide form a kinome annotation resource on LECA level for newly sequenced eukaryotic species. The first eukaryotic kinome tree is thus relevant both from an evolutionary, methodical and functional perspective.

## Methods

### Data collection

#### Eukaryotic proteome dataset

In order to collect ePK domains, 94 proteomes were carefully selected from available sequenced eukaryotic genomes. This proteome dataset was compiled as described earlier^49^ but with a slightly different, more diverse set of species. Four fungal species from the earlier dataset were removed while eight new species were added. The proteome dataset contains 1,538,389 proteins in total and is available (see **Data availability**). Details of the selected proteomes can be found in Supplementary Table 3. To each protein in the dataset, a unique protein identifier was assigned that is composed of four letters and six numbers. The four letters combine the first letter of the genus name with the first three letters of the species name. A bifurcating species tree of the species in the eukaryotic proteome dataset was assembled manually. This species tree was rooted between Amorphea and Bikonta^19^ and used as input for the LECA clade annotation pipeline.

#### Kinase domain dataset

From the eukaryotic proteome dataset, kinase domains were selected using PFAM models. All HMM profiles of PFAM-A version 31.0 were downloaded from the PFAM database^50^. These PFAM-A models were used in an HMMSCAN search (HMMER, http://hmmer.org, version 3.0) against the eukaryotic proteome dataset (bit score threshold: PFAM Trusted Cutoff). If a particular PFAM-A model hit the same protein sequence multiple times, domain bit scores of non-overlapping hits were summed (a maximum overlap of 30 amino acids was allowed). For each sequence, the best hitting non-overlapping PFAM models were determined based on these modified bit scores. Sequences that were best hit by PFAM models Pkinase and Pkinase_Tyr were collected, and kinase domains were excised based on envelope coordinates. In total, 28,249 Pkinase and 8,226 Pkinase_Tyr kinase domains were collected. The resulting fasta file with 36,475 kinase domains (Supplementary Data 1) was used as input for the LECA clade annotation pipeline.

#### LECA clade annotation pipeline

The LECA clade annotation pipeline (summarized in Fig. 1) is a Snakemake workflow that consists of a collection of rules in a Snakefile^51^. The rules in the Snakefile describe how to create output files from input files. The rules of the LECA clade annotation pipeline Snakefile and their interrelationships are illustrated in a Directed Acyclic Graph (DAG) in Supplementary Fig. 3. To run the LECA clade annotation pipeline, in addition to the Snakefile several data files, scripts and programmes are required. Snakefile, data files and scripts are available (see **Data availability**) while the programmes that were used are listed in the Snakefile. The LECA clade annotation pipeline itself was executed with Snakemake (version 3.11.2). Below, the different steps of the LECA clade annotation pipeline are described, and corresponding Snakefile rules are given in italics.

#### BBH selection

In order to enable generating a eukaryotic kinome tree, the LECA clade annotation pipeline starts with reducing the kinase domain dataset. The kinase domain dataset was reduced by selecting Bi-directional Best Hits (BBHs) between eukaryotic supergroups. For selecting BBHs, the fasta file with all 36,475 kinase domains (Supplementary Data 1) was divided into separate per species fasta files. These per species kinase fasta files were used in an all species versus all species BLASTp^52^ (version 2.3.0) run (rule *run_species_vs_species_blast)*. Based on combining all these BLAST searches, BBHs between eukaryotic supergroups were selected (rules *select_bbh_ids* and *collect_bbh_sequences*). Two different sets of BBHs were compiled. These two different datasets served three purposes: (1) having two different datasets enabled to check results for consistency, (2) a phylogenetic tree with a smaller number of BBHs was easier to annotate with LECA clades while (3) a phylogenetic tree with a larger number of BBHs allowed generating LECA clade HMM profiles with more sequence diversity. The first dataset, the two-supergroups-BBHs dataset, consists of 596 sequences that are BBHs between the two supra-supergroups Amorphea and Bikonta (Supplementary Data 2). The second dataset, the five-supergroups-BBHs dataset, consists of 1,738 sequences that are BBHs between five supergroups that are subsets of either Amorphea ((1) Opisthokonta + Apusozoa and (2) Amoebozoa) or Bikonta ((3) Archaeplastida + Cryptista, (4) SAR (Stramenopiles, Alevolata and Rhizaria) + Haptophyta and (5) Excavata) (Supplementary Data 6). The two-supergroups-BBHs dataset is a subset of the five-supergroups-BBHs dataset.

#### Phylogenetic tree generation

Both the two-supergroups-BBHs dataset and the five-supergroups-BBHs dataset were used to generate a phylogenetic tree. First, both datasets were aligned using mafft-einsi^53^ (version 7.127) (rule *run_mafft*). Positions in the alignments that did not have a gap score of at least 0.25 were removed with trimAl^54^ (version 1.2rev59) (rule *run_trim_al*). The resulting two-supergroups-BBHs alignment is 263 positions long while the five-supergroups-BBHs alignment contains 261 positions. Alignments were converted to Phylip format (rule *converse_alignment_to_phylip*) and made suitable for RAxML input (rule *prepare_headers_for_raxml*) by changing some characters in the sequence headers (Supplementary Data 3 and 7). Secondly, with RAxML^55^ (version 8.1.1) two maximum likelihood phylogenetic trees were generated using 100 rapid bootstraps, the GAMMA model of rate heterogeneity and an automatically determined best protein substitution model (rule *run_raxml*). For both trees, the best protein substitution model was LG. Annotated versions of the Newick trees and accessory files (see ***Newick file annotation***) are available in Supplementary Data 4, 5, 8 and 9.

#### LECA clade annotation with Notung

The kinase domain trees were analysed to determine clades (a.k.a. Orthologous Groups (OGs)) that were putatively one kinase in the LECA. As a first step to annotate the trees with these LECA clades or LECA OGs, the trees were rearranged with the gene tree-species tree reconciliation software package Notung^56^ (version 2.8.1.6-beta). Notung annotates duplication and speciation nodes in rearranged gene trees. As gene trees are imperfect, Notung was run with two different rearrangement cut-offs. This allowed flexibility in identifying LECA clades.

Leaves of both the two- and five-supergroups-BBHs trees were prepared for use within Notung by adding supergroup postfixes and species prefixes to leaf names (rules *add_supergroups_to_leaf_names* and *add_species_prefixes_for_notung*). The trees were also midpoint rooted with ETE^57^ (version 3.0.0b29) before running Notung because the default implicit rooting from RAxML could hamper LECA clade annotation at the outer edge of the trees. The two- and five-supergroups-BBHs trees were then rearranged with Notung according to the bifurcating species tree (rule *run_notung*) in the following manner: both trees were rearranged twice, once with bootstrap values below 50 allowed to be rearranged and once with bootstrap values below 70 allowed to be rearranged. The rearranged trees were stripped of species prefixes because the prefixes were redundant after running Notung (rule *remove_species_prefixes*).

In the four rearranged Notung trees, LECA clades were determined using a custom script that started with pre-LECA duplication nodes (rule *determine_notung_ogs*). Pre-LECA duplication nodes represent gene duplications that preceded all species and thus occurred before the LECA originated. They were parsed from a list of duplication nodes that Notung offers as output. In the rearranged phylogenetic trees, nodes that are children of pre-LECA duplication nodes but are not pre-LECA duplication nodes themselves were assessed as potential LECA speciation nodes. Potential LECA speciation nodes with at least one Amorphea and one Bikonta sequence among their children were defined as definitive LECA speciation nodes. All sequences descending from a definitive LECA speciation node were then defined as forming a single LECA clade. The reliability of a LECA clade that was annotated in a rearranged Notung tree was determined by evaluating the corresponding original RAxML tree. A LECA clade was regarded reliable if all sequences belonging to it also formed a single clade in the original RAxML tree. LECA clades that were not monophyletic in the original RAxML trees were labelled dubious and removed (rule *determine_dubious_notung_ogs*).

Because two different bootstrap cut-offs (50 and 70) were used to generate rearranged Notung trees, two different sets of LECA clades were available per original RAxML tree. For each RAxML tree, the two sets of LECA clades were combined into one set that attempted to annotate as many tree leaves as possible (rule *combine_notung_ogs*). The rationale behind maximizing the number of annotated leaves is that each present-day kinase likely originates from a LECA kinase clade. Preferably, LECA clades based on the most stringent rearrangement cutoff 70 (70-clades) were used for leaf annotation because 70-clades are better supported than clades based on rearrangement cutoff 50 (50-clades). However, 70-clades that were labelled as dubious or had bootstrap support below 50 were not used in the two combined sets of LECA clades. If possible, they were replaced by one or more 50-clades. Only non-dubious 50-clades with minimal bootstrap support of 50 could serve as a replacement. For the two-supergroups-BBHs tree, the combined set of LECA clades based on Notung consists of 117 LECA clades that annotate 535 of the 596 leaves.

For the five-supergroups-BBHs tree, the combined set consists of 113 LECA clades that annotate 1,283 of the 1,738 leaves.

#### LECA clade annotation with HMMER

In a second tree annotation step, LECA clades annotated with Notung were expanded with not yet annotated tree leaves using HMMER. After annotating putative LECA clades with Notung, several sequences in the trees were not annotated, even though they resided close by annotated Notung clades. These leaves were prevented from being part of a Notung LECA clade by bootstrap values or inconsistencies between gene tree and species tree. To still annotate these leaves with existing Notung LECA clades, the two combined sets of Notung LECA clades were projected on the two BBH sets in two consecutive rounds of HMMER searches.

Notung LECA clade HMM profiles for the first HMMER search against BBHs were generated as follows. For both the two- and five-supergroups-BBHs, BBH sequences that were annotated as part of a Notung LECA clade were collected for each Notung LECA clade in a fasta file (rule *distribute_ogs_1*). Each Notung LECA clade fasta file was aligned with mafft-einsi, and the alignment was used to generate a HMMER3 profile. All two-supergroups-BBHs Notung LECA clade HMMER3 profiles were used for a HMMER search against the two-supergroups-BBH sequences (rule *run_hmmer_search_bbhs_1*). The five-supergroups-BBHs Notung LECA clade HMMER3 profiles were used for a similar search against the five-supergroups-BBH sequences. A list of top two best hitting Notung LECA clade HMM profiles was generated for both the two- and five-supergroups-BBHs (rule *determine_hmmer_ogs_bbhs_1*). Only BBHs with a bit score difference of minimal 10 between top two hits and at least one bit score of minimal 30 were listed. Leaves of the two- and five-supergroups-BBH trees that were not yet annotated with Notung were then annotated with the Notung LECA clade that was their best HMMER hit in the top two list (rule *add_hmmer_ogs_bbhs_1*). Leaf annotation with the best Notung LECA clade HMMER hit was only completed provided the Notung LECA clade leaves and the leaf best hit by the associated Notung LECA clade HMMER profile were monophyletic in the rooted RAxML tree. In the first round of HMMER annotation, 42 leaves of the two-supergroups-BBHs tree and 167 leaves of the five-supergroups-BBHs tree were annotated with HMMER in addition to the Notung annotation.

The expansion of Notung LECA clades with leaves annotated with HMMER allowed generating more sensitive HMMER profiles. These more sensitive HMMER profiles for the second HMMER search against BBHs were generated as follows. BBH sequences that were annotated with Notung or HMMER as forming a single LECA clade were combined, aligned with mafft-einsi and subsequently a Notung-HMMER LECA clade HMMER3 profile was generated (rule *distribute_ogs_2*). All two-supergroups-BBHs Notung-HMMER LECA clade HMMER3 profiles were combined into one set, and the same was done for all five-supergroups-BBHs Notung-HMMER LECA clade HMMER3 profiles. These two sets of HMMER profiles were each used for a HMMER search against both the two- and five-supergroups-BBH sequences (rule *run_hmmer_search_bbhs_2*). The HMMER searches with HMMER profiles corresponding to their source tree were used for further leaf annotation (e.g. two-supergroups-BBHs Notung-HMMER LECA clade profiles against two-supergroups-BBHs). The HMMER searches with HMMER profiles derived from the other tree were later in the pipeline used for mapping leaf annotation between the two- and five-supergroups-BBH trees (e.g. two-supergroups-BBHs Notung-HMMER LECA clade profiles against five-supergroups-BBHs). For each of the four HMMER searches a list of each BBHs top two best hitting Notung-HMMER LECA clade HMMER profiles was generated (rule *determine_hmmer_ogs_bbhs_2*). This was done under the same conditions as described for the first round of HMMER annotation. Leaves of the two- and five-supergroups-BBH trees that were not yet annotated with Notung or the first round of HMMER annotation were then annotated with the best hitting Notung-HMMER LECA clade from their respective Notung-HMMER LECA clade set (rule *add_hmmer_ogs_bbhs_2*). Leaf annotation was again only completed provided the Notung-HMMER LECA clade leaves and the leaf best hit by the associated Notung-HMMER LECA clade HMMER profile were monophyletic in the rooted RAxML tree. In this second round of HMMER annotation, three leaves of the two-supergroups-BBHs tree and 49 leaves of the five-supergroups-BBHs tree were annotated on top of the existing annotation. In total, 580 of the 596 leaves of the two-supergroups-BBHs tree and 1,499 of the 1,738 leaves of the five-supergroups-BBHs tree were annotated. Notung-HMMER LECA clade sequences and sequences newly annotated with the same LECA clade in the second round of HMMER searches were combined in per LECA clade fasta files (rule *distribute_ogs_3*).

#### Combination LECA clades two- and five-supergroups-BBHs trees

In a third step to annotate the trees with LECA clades, Notung-HMMER LECA clades of the two- and five-supergroups-BBHs trees were combined into one overarching set. To do this, first for each tree a list with leaves was generated that per leaf provides the two Notung-HMMER LECA clades to which the leaf is annotated in respectively the two- and five-supergroups-BBHs trees (rule *map_ogs*). This list was subsequently used to extract which Notung-HMMER LECA clade in the two-supergroups-BBHs tree corresponds to which Notung-HMMER LECA clade in the five-supergroups-BBHs tree and vice versa. If leaves of a Notung-HMMER LECA clade in one of the two trees were distributed over multiple Notung-HMMER LECA clades in the other tree, these multiple LECA clades were merged into a single LECA clade (rule *merge_ogs*) (Supplementary Table 4). After merging, the number of Notung-HMMER LECA clades annotated in the two-supergroups-BBHs tree decreased from 117 to 110.

Based on the mapping of Notung-HMMER LECA clades between the two- and five-supergroups-BBHs trees and the merged LECA clades, a new set of combined LECA clades was generated (rule *combine_ogs*). The 110 Notung-HMMER LECA clades that were annotated in the two-supergroups-BBHs tree and the 113 Notung-HMMER LECA clades that were annotated in the five-supergroups-BBHs tree formed together 118 unique Notung-HMMER LECA clades. Eight Notung-HMMER LECA clades were absent in the two-supergroups-BBHs tree, and five Notung-HMMER LECA clades were not automatically annotated in the five-supergroups-BBHs tree (Supplementary Table 5). Per combined LECA clade, sequences of the Notung-HMMER LECA clade(s) that form the combined LECA clade were collected, aligned with mafft-einsi and used to generate a HMMER3 profile (rule *combine_og_profiles*). A modified version of these HMMER3 profiles (see ***Manual Annotation***) is available in Supplementary Data 10.

#### Domain assignment

The combined set of 118 unique Notung-HMMER LECA clades was used to assign the initial 36,475 kinase domains to a LECA clade. In order to do this, the combined LECA clade HMMER3 profiles were run against the complete kinase domain dataset (rule *run_hmmer_search_all_4*). The results of this HMMER run were used to divide kinase domains over three lists (rule *determine_hmmer_ogs_all_4*): (1) assigned kinase domains with a bit score difference of minimal 10 between top two LECA clade hits and at least one LECA clade hit with a bit score of minimal 30, (2) difficult-to-assign kinase domains with a bit score difference below 10 between top two LECA clade hits and (3) unassigned kinase domains with only LECA clade hits with a bit score below 30. Modified versions of these lists (see ***Manual Annotation***) are available as Supplementary Data 11-13. The assigned kinase domains from the first list were used to generate a matrix that for each species in the eukaryotic proteome dataset and for each combined LECA clade provides how many sequences were hit. An adjusted version of this matrix (Supplementary Data 14) forms the basis for Fig. 3 to 6. Furthermore, the assigned kinase domains were classified per eukaryotic supergroup to determine the number of eukaryotic supergroups hit by each LECA clade. A supergroup was only counted if kinases from minimal two species of that supergroup were hit by a particular combined LECA clade. Per species, the percentage of assigned kinase domains was also determined (rule *determine_assignment_percentages*) (Supplementary Table 2).

#### LECA clade categorization

The reliability of the set of 118 combined LECA clades was evaluated using information directly and indirectly available in the LECA clade annotation pipeline. The combined LECA clades were therefore classified in categories with respect to the amount of support they have from the two different trees and domain assignment. Per combined LECA clade, the following information was listed (rule *add_hmmer_supergroups*): (1) presence/absence in both RAxML trees, (2) number of eukaryotic supergroups among assigned kinases using the counting mode of the supergroup matrix described under ***Domain assignment***, (3) bootstrap support in both RAxML trees, and (4) correspondence to multiple Notung-HMMER LECA clades in one of the RAxML trees. Based on this information, LECA clades were labelled with four categories (rule *add_categories*): (1) combined LECA clades that are annotated only in one of the two trees (indicated with *), (2) combined LECA clades to which domains from less than two eukaryotic supergroups were assigned with HMMER (indicated with **), (3) combined LECA clades that in neither of the RAxML trees have bootstrap support of minimal 70 (indicated with ***) and (4) combined LECA clades that in one of the trees are split in multiple Notung-HMMER LECA clades (indicated with %). A fifth category was added later upon manual annotation (see ***Manual annotation***). Not all LECA clades are labelled with any of the categories while LECA clades can also be labelled with multiple categories at once.

#### LECA clade naming

The 118 combined LECA clades were named in order to distinguish them better and easily link them with functional information (rule *make_og_name_table_and_list*). They were named by their best hit in the well-studied species human, baker’s yeast or *A. thaliana.* Preferably, human kinases were used for naming, but if no human kinase was assigned to a LECA clade, baker’s yeast or *A. thaliana* names were used. If hits from all three species were absent, a LECA clade was indicated with its combined LECA clade number. Furthermore, all kinases from human, baker’s yeast and *A. thaliana* that were assigned to a LECA clade were listed in a table (Supplementary Table 1). Per LECA clade, hits from these species were displayed in descending bit score order. LECA clades in the table were extended with categories (rule *add_og_categories_to_table*). LECA clade names were also added to earlier generated files (rules *add_og_names_to_list* and *add_og_names_to_matrix*).

#### Newick file annotation

To browse through the eukaryotic kinome tree and LECA clade annotation easily at once, leaf names of the Newick files of the two- and five-supergroups-BBHs trees were extended with LECA clade annotation. Leaf names had already been extended with supergroup names before (rule *add_supergroups_to_leaf_names*). If leaves were annotated with Notung of HMMER, they were extended with a Notung LECA clade (abbreviated as nOG) (rule *add_notung_ogs_to_leaf_names*) or HMMER LECA clade (abbreviated as hOG) (rules *add_hmmer_ogs_to_leaf_names_bbhs_1* and *add_hmmer_ogs_to_leaf_names_bbhs_2*). Subsequently, leaves that participated in a combined LECA clade were extended with this combined LECA clade (abbreviated as cOG) including both its number, categories and name (rule *add_combined_ogs_to_leaf_names*). Leaves from combined LECA clades that include manually annotated leaves (see ***Manual annotation***) were denoted as mOG instead of cOG. Finally, to all leaves the top two best hitting combined LECA clades were added together with their bit scores (rule *add_combined_og_hmmer_hits_to_leaf_names*). For visualising the trees with iTOL^58^, nodes that are the common ancestor of leaves that form a LECA clade were named with this LECA clade (rule *add_ogs_to_nodes*). Furthermore, to automatically collapse the LECA clades in iTOL, a ‘collapse file’ was generated. Annotated Newick files of the two- and five-supergroups-BBHs trees (Supplementary Data 4 and 8) and their collapse files (Supplementary Data 5 and 9) form the basis for Fig. 2 and Supplementary Fig. 1.

#### Manual annotation

When inspecting the annotated Newick trees, the automatic LECA clade annotation displayed room for manual improvement. Manual annotation was performed in the following cases: (1) to split merged LECA clades (indicated with %) if there were good reasons to believe that they are indeed multiple LECA clades, (2) to merge LECA clades if there were good reasons to believe that they are indeed one LECA clade or (3) to annotate not yet annotated leaves. Manual annotation was done by partially re-executing the LECA clade annotation pipeline (starting with rule *run_hmmer_search_all_4*) after copying and manually changing files including the HMMER3 profiles of combined LECA clades (rule *copy_lists_notung_hmmer_hmmer_ogs* and the description of manual runs in the Snakefile). Manual annotation occurred in two rounds, with annotated Newick trees of the first manual run serving to determine the next step in the second manual run. The LECA clades that include manually annotated leaves and the reasoning that resulted in their manual annotation are described in Supplementary Table 6. In total, 16 LECA clades were fully or partially based on manually annotated leaves.

The total number of combined LECA clades did not change after manual annotation and remained 118. But the number of annotated combined LECA clades in the two-supergroups-BBHs tree increased from 110 to 113 and the number of annotated combined LECA clades in the five-supergroups-BBHs tree decreased from 113 to 111 (Supplementary Table 6).

Manually annotated LECA clades form a fifth category (see ***LECA clade categorization***) that is indicated with #. In the Newick trees, their leaves are indicated with mOG instead of cOG. In the two-supergroups-BBHs tree, 17 of the 596 leaves were manually annotated resulting in the final annotation of 593 leaves. In the five-supergroups-BBHs tree, 264 of the 1,738 leaves were manually annotated resulting in the final annotation of 1,585 leaves.

## Figure generation

Fig. 1 was produced with Lucidchart, https://www.lucidchart.com. Fig. 2 was produced with iTOL^58^ (version 4.3) based on Supplementary Data 4 and 5. Fig. 3 to 6 were produced with R^59^ (version 3.4.4) based on Supplementary Data 14 and 16, using R packages APE^60^ and gplots^61^. All figures were adjusted with Inkscape, http://inkscape.org.

## Data availability

The eukaryotic proteome dataset is available at https://bioinformatics.bio.uu.nl/snel/support/eukaryotic_proteome_dataset. The entire LECA clade annotation pipeline, including all input data and output files, is available at https://bioinformatics.bio.uu.nl/snel/support/LECA_clade_annotation_pipeline. A selection of output files is also available as Supplementary Data.

## Code availability

The computational code of the LECA clade annotation pipeline is together with the data available at https://bioinformatics.bio.uu.nl/snel/support/LECA_clade_annotation_pipeline.

## Acknowledgements

We thank Sebastiaan Broekema for carrying out a pilot project on the usability of the Scrollsaw method for the eukaryotic kinome, John van Dam for his contribution to compiling the eukaryotic proteome dataset and members of the Snel lab for critical reading and helpful discussion on the manuscript. We thank Ivica Letunic for support with iTOL. This research was financially supported by the Netherlands Organization for Scientific Research (NWO) grant 016.160.638 Vici.

## Author information

### Contributions

B.S. and L.M.W. designed the research. L.M.W. performed the research and analysed the data. B.S. and L.M.W. wrote the manuscript.

## Supplementary material

Supplementary Information:

Description of Supplementary Tables

Description of Supplementary Data

Supplementary Results

Supplementary References

Supplementary Fig. 1-3

Supplementary Tables 1-16

Supplementary Data 1-16

