## Supplementary Information for "The first eukaryotic kinome tree illuminates the dynamic history of present-day kinases"

### **Supplementary information for** Van Wijk and Snel**, The first eukaryotic kinome tree illuminates the dynamic history of present-day kinases**

**Supplementary Information** **1**
Description of Supplementary Tables 1
Description of Supplementary Data 4
Supplementary Results 5
Supplementary References 13

#### Description of Supplementary Tables

**Supplementary Table 1: LECA clade assignment of ePK domains from *Homo sapiens*, *Saccharomyces cerevisiae* and *Arabidopsis thaliana*.** Per species and LECA kinase clade, domains are ordered in decreasing bit score order. This table is based on the output of rules *make_og_name_table_and_list* and *add_og_categories_to_table*.

**Supplementary Table 2: LECA clade assignment of ePK domains from 94 present-day eukaryotes.** Per species, numbers and percentages of assigned domains, difficult-to-assign domains, domains that are only hit below the assignment bit score cut off and domains not hit by any LECA kinase clade are given. This table is based on the output of rule *determine_assignment_percentages*.

**Supplementary Table 3: Sources of the 94 eukaryotic proteomes in the eukaryotic proteome dataset.**

**Supplementary Table 4: Merged LECA kinase clades.**

LECA kinase clades are listed that in the original annotation of the two- and five-supergroups-BBHs trees with Notung-HMMER LECA clades were annotated as one LECA kinase clade in one tree and as two LECA kinase clades in the other tree.

**Supplementary Table 5: LECA kinase clades that are not annotated in both BBHs trees.**

LECA kinase clades are listed that in one of the two BBHs trees are absent, not automatically annotated or not annotated after manual splitting. Some of the listed LECA kinase clades are in the final LECA kinase clade annotation manually annotated or manually merged with other LECA kinase clades (Supplementary Table 6).

**Supplementary Table 6: Manually annotated LECA kinase clades.** LECA kinase clade annotation adjustments are given per manual annotation round and per BBH tree, and considerations involved in manual annotation are described.

**Supplementary Table 7: Well-supported pre-LECA kinase clades.** Pre-LECA kinase clades that exist of more than two LECA kinase clades are indicated with a grey background. Pre-LECA kinase clades that are parents of other pre-LECA kinase clades are indicated in bold. Pre-LECA kinase clades that are children of other pre-LECA kinase clades are indicated in italics. Bootstrap support below 70 is also indicated in italics. Numbers of pre-LECA kinase clades that are well-supported in both trees are indicated in bold. If these pre-LECA kinase clades do not agree with the human kinome tree, the disagreement is also indicated in bold. For pre-LECA kinase clades that are retrieved in both trees but disagree with the human kinome tree, rearrangements to the human kinome tree are proposed.

**Supplementary Table 8: Moderately supported pre-LECA kinase clades.** Pre-LECA kinase clades that exist of more than two LECA kinase clades are indicated with a grey background. Pre-LECA kinase clades that are parents of other pre-LECA kinase clades are indicated in bold. Pre-LECA kinase clades that are children of other pre-LECA kinase clades are indicated in italics. Bootstrap support below 50 is also indicated in italics. Numbers of pre-LECA kinase clades that are moderately supported in both trees are indicated in bold. If these pre-LECA kinase clades do not agree with the human kinome tree, the disagreement is also indicated in bold and rearrangements to the human kinome tree are proposed.

**Supplementary Table 9: Basal ePKs[3] and corresponding LECA kinase clades.**

**Supplementary Table 10: Classification of LECA kinase clades into new and already known LECA kinase clades.** If a basal ePK corresponds to multiple LECA kinase clades (Supplementary Table 9), these multiple LECA kinase clades are only once classified as already known.

**Supplementary Table 11: Human kinases unique to the eukaryotic proteome dataset.**

**Supplementary Table 12: Human kinases unique to the human kinome tree.**

**Supplementary Table 13: EPK superfamily support and composition.**

**Supplementary Table 14: LECA clade assignment of 'Other' (Unaffiliated) kinases from the human kinome tree.** 'Other' kinases that are classified as 'Other' in Supplementary Table 1 of the Human Kinome Paper[1] but are not located on a grey branch in the human kinome tree (Supplementary Fig. 2) are indicated in italics. Kinases that form well-supported pre-LECA kinase clades are located beneath each other and have the same background colour. Kinases that form moderately supported pre-LECA kinase clades are located beneath each other and are surrounded by a dashed line. LECA kinase clades that do not exist exclusively of 'Other' kinases are indicated in italics. LECA kinase clades that are not monophyletic in the human kinome tree are indicated in bold.

**Supplementary Table 15: Polyphyletic LECA kinase clades in the human kinome tree and proposed rearrangements.**

**Supplementary Table 16: Unassigned human kinases.** For each unassigned human kinase, the top two best hitting LECA kinase clades are given. Top two LECA kinase clades and kinase superfamilies that are in line with the assignment of kinases that in the human kinome tree are monophyletic with the unassigned kinase are indicated in bold. For remaining kinases, ePK superfamilies are indicated in italics if top two best hitting LECA kinase clades are from the same ePK superfamily. This table is based on a subset of Supplementary Data 12.

#### Description of Supplementary Data

**Supplementary Data 1: Initial fasta file with 36,475 ePK domains.** The file is compiled of sequences best hit by PFAM models Pkinase (28,249 domains) and Pkinase_Tyr (8,226 domains).

**Supplementary Data 2: Fasta file with 596 BBHs between two eukaryotic supergroups.**

**Supplementary Data 3: Two-supergroups-BBHs alignment.** The alignment is trimmed and prepared for RAxML.

**Supplementary Data 4: Two-supergroups-BBHs Newick tree.** The tree is annotated with the final set of 118 LECA kinase clades. Leaf names are extended as described in the Methods.

**Supplementary Data 5: Collapse file for the two-supergroups-BBHs Newick tree.** This file can be uploaded in iTOL together with Supplementary Data 4.

**Supplementary Data 6: Fasta file with 1,738 BBHs between five eukaryotic supergroups.**

**Supplementary Data 7: Five-supergroups-BBHs alignment.** The alignment is trimmed and prepared for RAxML.

**Supplementary Data 8: Five-supergroups-BBHs Newick tree.** The tree is annotated with the final set of 118 LECA kinase clades. Leaf names are extended as described in the Methods.

**Supplementary Data 9: Collapse file for the five-supergroups-BBHs Newick tree.** This file can be uploaded in iTOL together with Supplementary Data 8.

**Supplementary Data 10: HMM profiles for all 118 LECA kinase clades.**

**Supplementary Data 11: List of ePKs that are assigned to a LECA kinase clade.** Per ePK, the top two best hitting LECA kinase clades and their associated bit scores are given.

**Supplementary Data 12: List of difficult-to-assign ePKs.** These ePKs were not assigned to a LECA kinase clade because of a bit score difference below 10 between top two hits. Per ePK, the top two best hitting LECA kinase clades and their associated bit scores are given.

**Supplementary Data 13: List of unassigned ePKs.** These ePKs were not assigned to a LECA kinase clade because none of the LECA kinase clade HMM profiles were hit with a minimal bit score of 30.

**Supplementary Data 14: Matrix with LECA kinase clade assignment for 28,893 ePK domains from 94 eukaryotes.** This matrix is the output of rule *determine_hmmer_ogs_all_4*, with names added by rule *add_og_names_to_matrix*. The matrix is manually extended with a column that classifies LECA kinase clades into ePK superfamilies.

**Supplementary Data 15: Fasta file with 130 NAF domains, including that of ePK TTRA004140.** This file is a collection of sequences best hit by PFAM model NAF. It is generated in the same way as the files with Pkinase and Pkinase_Tyr domains that together form Supplementary Data 1.

**Supplementary Data 16: R script that generates Fig. 3 – 6 based on Supplementary Data 14.**

#### Supplementary Results

In the Supplementary Results, claims described in the main Results will be further substantiated.

**Well-supported and moderately supported pre-LECA kinase clades**

For elucidating deep relations between LECA kinase clades, internal support in the eukaryotic kinome tree is important. As stated in the main Results, LECA kinase clades are well supported, but support for pre-LECA kinase clades is much lower. In the two-supergroups-BBHs tree, 112 pre-LECA kinase clades are found (Fig. 2) while in the five-supergroups-BBHs tree, 110 pre-LECA kinase clades are present (Supplementary Fig. 1). A total of 34 pre-LECA kinase clades is well-supported (minimal bootstrap support 70) in at least one of both trees, including 12 pre-LECA kinase clades that consist of three or more LECA kinase clades (Supplementary Table 7). The number of pre-LECA kinase clades that are well-supported in both trees is 23, including seven of the pre-LECA kinase clades that consist of three or more LECA kinase clades. For another six of the 34 pre-LECA kinase clades that are well-supported in at least one tree, bootstrap support in the other tree is between 50 and 70. In addition to the 34 well-supported pre-LECA kinase clades, eight pre-LECA kinase clades are moderately supported (minimal bootstrap support 50) in at least one tree and retrieved in both trees (Supplementary Table 8). Although their support is not high, the consistency of these pre-LECA kinase clades in both trees suggest they may be correct. Seven of them include three or more LECA kinase clades. The well-supported and moderately supported deeper relationships between LECA kinase clades provide a starting point to transfer functional information between paralogous LECA kinase clades. They could also elucidate the order of emergence of eukaryotic cellular innovations in which these LECA kinase clades play a role.

**Comparison between LECA kinase clades and an earlier LECA kinase number estimate**

By comparing the LECA kinase clades in the eukaryotic kinome tree to an earlier LECA kinome estimate and by evaluating different sources of support, a nuanced estimation of the LECA ePK complement was possible. Our conservative estimate of 92 LECA ePKs is more than a third more than the largest previous estimate of 68[3]. These estimates of LECA ePKs were calculated as follows.

The largest previous estimate of 68 LECA ePKs is based on a set of 88 basal kinases that includes 70 ePKs[3] (Supplementary Table 9). Two basal ePKs from this study (CAMK:CDPK and CK1:CK1:CK1y) did not meet the LECA definition of being present in both Amorphea and Bikonta, as they are absent in Amorphea. When these two are excluded, 68 basal ePKs agree with a LECA definition of presence in both Amorphea and Bikonta. Out of these 68 basal ePKs, 66 ePKs were found back in the 118

LECA kinase clades that are annotated in the eukaryotic kinome tree. One basal ePK, Other:Haspin was missing among the LECA kinase clades because it was absent in the initial set of 36,475 ePKs (see also ***Dataset overlap*** below). The second 'missing' basal ePK is caused by two basal ePKs, CMGC:CDK:CDC2 and CMGC:CDK:CDK5, that form one LECA kinase clade in the eukaryotic kinome tree: OG074 CDK3. Based on the topologies of the two- and five-supergroups-BBHs trees, it cannot be excluded that OG074 CDK3 consists of two nested LECA kinase clades.

The 66 basal ePKs that were found among the 118 LECA kinase clades correspond to 72 LECA kinase clades. The six additional LECA kinase clades originate from three basal ePKs (CMGC:CDKL, Other:SCY1, Other:ULK) that each correspond to two LECA kinase clades (OG076 CDKL2 and OG077 CDKL5, OG036 SCYL1 and OG037 SCYL2, OG063 ULK1 and OG089 ULK4) and one basal ePK (TKL) that corresponds to four LECA kinase clades (OG110 LIMK1, OG111 TESK2, OG112 IRAK4 and OG114*/*** ZAK) (Supplementary Tables 9 and 10). Apart from these six additional LECA kinase clades compared to the 66 basal ePKs, another 46 completely new LECA kinase clades were found (Supplementary Table 10). Therefore, in total, 52 new LECA kinase clades were found in the eukaryotic kinome tree. Thirty-one of the 52 new LECA kinase clades are annotated in both trees, well-supported in at least one tree (minimal bootstrap support 70) and kinases from minimal two eukaryotic supergroups are assigned to them. Based on these numbers, a conservative estimate of LECA ePKs would be 98: 66 LECA kinase clades that overlap with earlier defined basal ePKs, plus Haspin, plus 31 new LECA kinase clades. However, also within the LECA kinase clades that overlap with basal ePKs six LECA kinase clades do not full-fill all three criteria of being annotated in both trees, being well-supported and being present in sufficient species. Subtracting them yields an even more conservative estimate of 92 kinases in the LECA.

#### Comparison between eukaryotic kinome tree and human kinome tree

No formal benchmark for kinase evolution exists but an insightful evaluation of the outcome of the pipeline that generated the eukaryotic kinome tree is provided by the human kinome tree (Supplementary Fig. 2). Hence the eukaryotic kinome tree was directly compared to the human kinome tree. In the main Results, the eukaryotic kinome tree and human kinome tree were stated to be highly consistent, although full agreement requires some adjustments to the human kinome tree. Here a detailed comparison between the two trees is provided concerning dataset overlap, superfamily classification and agreement between (pre-)LECA kinase clades and clades in the human kinome tree. Furthermore, the potential origin of human kinases that are unassigned in the eukaryotic kinome tree is described.

##### *Dataset overlap*

The initial set of 36,475 ePK domains that were used to generate the eukaryotic kinome tree includes 493 human ePK domains stemming from 480 proteins. The 13 proteins with double ePK domains in this dataset are TYK, JAK1, JAK2 and JAK3 (both domains TK/TKL superfamily), RSK1, RSK2, RSK3, RSK4, MSK1 and MSK2 (one domain CAMK, one domain AGC superfamily), GCN2 (one domain STE superfamily, one domain Unaffiliated), SPEG and OBSCN (both domains CAMK superfamily). The dataset that underlies the human kinome tree contains 518 kinases, including 478 ePKs[1]. Because this dataset of 478 ePKs contains the 13 double ePK domain proteins, 491 ePK domains are present in the human kinome tree. The overlap between the two datasets is 485 ePK domains. The eukaryotic kinome tree dataset contains eight unique ePK domains (Supplementary Table 11), while the human kinome tree dataset includes six unique human ePK domains (Supplementary Table 12). Thus the human part of the eukaryotic kinome tree dataset and the human kinome tree dataset overlap for no less than 97 per cent despite different data collection approaches.

***Superfamily classification***

The human kinome tree is classified into seven ePK superfamilies and a remaining class of 'Other' ePKs[1]. In this paper, 'Other' ePKs are referred to as Unaffiliated ePKs. The eukaryotic kinome tree is classified into the Unaffiliated ePKs and six ePK superfamilies: in the eukaryotic kinome tree, the TK and TKL superfamilies are nested and thus combined into a single TK/TKL superfamily. Within this TK/TKL superfamily,  the only LECA kinase clade to which human TK kinases are assigned is OG116 BTK.

The superfamily classification of human ePKs can be found both in the human kinome tree (Supplementary Fig. 2) and in Supplementary Table 1 of the Human Kinome Paper[1]. However, these classifications slightly differ. For the comparison between eukaryotic and human kinome tree, the classification in Supplementary Table 1 of the Human Kinome Paper is primarily followed. The classification in Supplementary Table 1 of the Human Kinome Paper differs from the human kinome tree in the following aspects:

- YANK1-3 are classified as AGC superfamily instead of Unaffiliated
- MOS, SGK496 and PBK are classified as Unaffiliated instead of TKL superfamily
- ANPA, ANPB, HSER, CYGD and CYF are classified as a separate, eighth ePK superfamily RGC instead of Unaffiliated. Because the RGC superfamily was not yet present in the LECA, in contrast to the other ePK superfamilies, the human kinome tree classification of RGC kinases as Unaffiliated was followed for comparison with the eukaryotic kinome tree.

The ePK superfamily structure of the human kinome tree is recapitulated in the eukaryotic kinome tree. All human kinases that belong to a single ePK superfamily in the human kinome tree are assigned to LECA kinase clades that form a single clade in the eukaryotic kinome tree (Fig. 2, Supplementary Fig. 1 and 2, Supplementary Table 1).  Only the TK and TKL superfamilies from the human kinome tree form an exception due to being nested in the eukaryotic kinome tree. All ePK superfamilies in the eukaryotic kinome tree are also well-supported (minimal bootstrap support 70) or moderately supported (minimal bootstrap support 50) in at least one of both BBHs trees (Supplementary Table 13).

In addition to recapitulating the ePK superfamily structure, the eukaryotic kinome tree clarified the relationships between several Unaffiliated human kinases and the ePK superfamilies. A majority of 56 of the 88 Unaffiliated kinases is assigned to 17 Unaffiliated (pre-)LECA kinase clades (Supplementary Table 14). However, 14 Unaffiliated human kinases are assigned to a LECA kinase clade residing within an ePK superfamily (Supplementary Tables 13 and 14).  Nine of these Unaffiliated kinases (SBK, SGK069, the five RGC kinases, MOS and SGK196) are assigned to LECA kinase clades to which human kinases are assigned that are not monophyletic in the human kinome tree (Supplementary Tables 1, 14 and 15, Supplementary Fig. 2). They likely exhibit high sequence divergence, at least in the human lineage. The remaining five Unaffiliated kinases form three LECA kinase clades (OG041 CAMKK2, OG078 CK2A1 and OG079 CDC7). The clustering of OG041 CAMKK2 within the CAMK superfamily is described in more detail in the main Results. LECA kinase clades OG078 CK2A1 and OG079 CDC7 cluster with the CMGC superfamily in the eukaryotic kinome tree. All human kinases assigned to OG078 CK2A1 and OG079 CDC7 were however already monophyletic with the CMGC superfamily in the human kinome tree despite not being classified as

CMGC kinases. Note that the classification of human OG078 CK2A1 members as CMGC kinases also has been proposed earlier[4,5].

***Agreement between LECA kinase clades and the human kinome tree***

Comparing the eukaryotic kinome tree to the human kinome tree reveals large consistency between LECA kinase clades in the eukaryotic kinome tree and clades in the human kinome tree. In human, kinases that descend from 82 different LECA kinase clades were retained (Supplementary Table 2). A vast majority of 71 of these 82 LECA kinase clades is monophyletic in the human kinome tree: human kinases assigned to such LECA kinase clades form single clades in the human kinome tree (Supplementary Fig. 2). The remaining 11 LECA kinase clades are polyphyletic in the human kinome tree: human kinases assigned to these LECA kinase clades do not form monophyletic clades in the human kinome tree (Supplementary Table 15). In some cases, further scrutiny of a polyphyletic LECA kinase clade is required first because it falls in a less trusted category (OG018A*/***/# PEK, OG114*/*** ZAK) or unassigned kinases close by in the human kinome tree suggest that LECA kinase clades might be missed or over split (OG063 ULK1, OG100# MST1, OG103 TTBK1 and OG114*/*** ZAK). For the other polyphyletic LECA kinase clades, OG006 NEK4, OG036 SCYL1, OG043# CAMK1D, OG046 MARK2, OG092# MAP3K3 and OG112 IRAK4, rearrangements to the human kinome tree are proposed (Supplementary Table 15).

***Agreement between pre-LECA kinase clades and the human kinome tree***

Pre-LECA kinase clades, consisting of two or more LECA kinase clades from the eukaryotic kinome tree, are also generally in agreement with the human kinome tree. Thirty-four pre-LECA kinase clades are well-supported (minimal bootstrap support 70) in at least one of both BBHs trees (Supplementary Table 7). A pre-LECA kinase clade is deemed suitable for comparison to the human kinome tree if the pre-LECA kinase clade is found in both BBHs trees and if human kinases are assigned to multiple of its constituent LECA kinase clades.  Twenty-six pre-LECA kinase clades fulfil these requirements.  A majority of 18 of the 26 pre-LECA kinase clades that were compared is in agreement with the human kinome tree. For the remaining eight pre-LECA kinase clades that disagree with the human kinome tree, rearrangements to the human kinome tree are proposed (Supplementary Table 7).  Another eight pre-LECA kinase clades to which human kinases are assigned were also compared to the human kinome tree (Supplementary Table 8). These pre-LECA kinase clades are moderately supported (minimal bootstrap support 50) in at least one BBHs tree but were retrieved in both trees. From these eight pre-LECA kinase clades, two are in agreement with the human kinome tree. The remaining six pre-LECA kinase clades disagree with the human kinome tree. Only for the two of them that are moderately supported in both trees, rearrangements to the human kinome tree are proposed (Supplementary Table 8).

***The origin of unassigned human kinases***

Not all human kinases were assigned to a LECA kinase clade in the eukaryotic kinome tree. Thirty-three human kinases remained unassigned due to a conservative assignment method that requires bit score differences above 10 between the top two hits. However, the top two hits of most of these 33 unassigned human kinases were consistent with their position in the human kinome tree. Eighteen of the unassigned human kinases form clades in the human kinome tree with other kinases that are all best hit by the same LECA kinase clade (Supplementary Table 16). For six more, their second best hit agrees with the assignment of human kinases that form a single clade with the unassigned kinase. For the nine remaining unassigned human kinases, there is no agreement in LECA kinase clades between unassigned and assigned kinases that form a single clade in the human kinome tree. However, in seven out of nine cases, the superfamily of the top two hits is the same, or both hits are Unaffiliated LECA kinase clades. This at least indicates the origin of these seven kinases at the ePK superfamily level.
