## Supplementary Figures 1-3 for "The first eukaryotic kinome tree illuminates the dynamic history of present-day kinases"

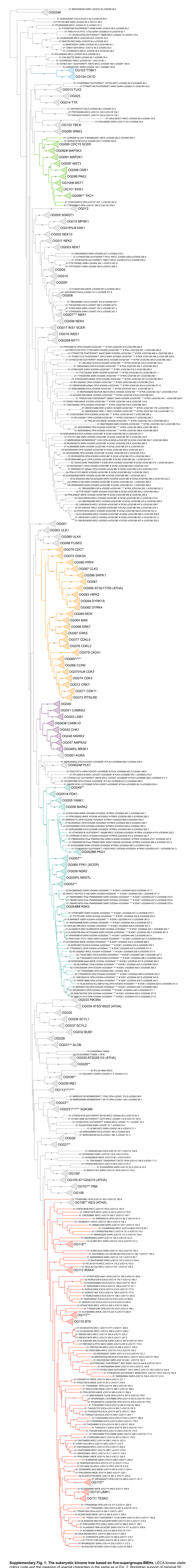


**Supplementary Fig. 1: The eukaryotic kinome tree based on five-supergroups-BBHs.** LECA kinase clade colour code and the meaning of special characters is the same as in Fig. 2. Bootstrap support of minimal 50 out of 100 is shown. This figure was produced with iTOL based on Supplementary Data 8 and 9 and adjusted with Inkscape.

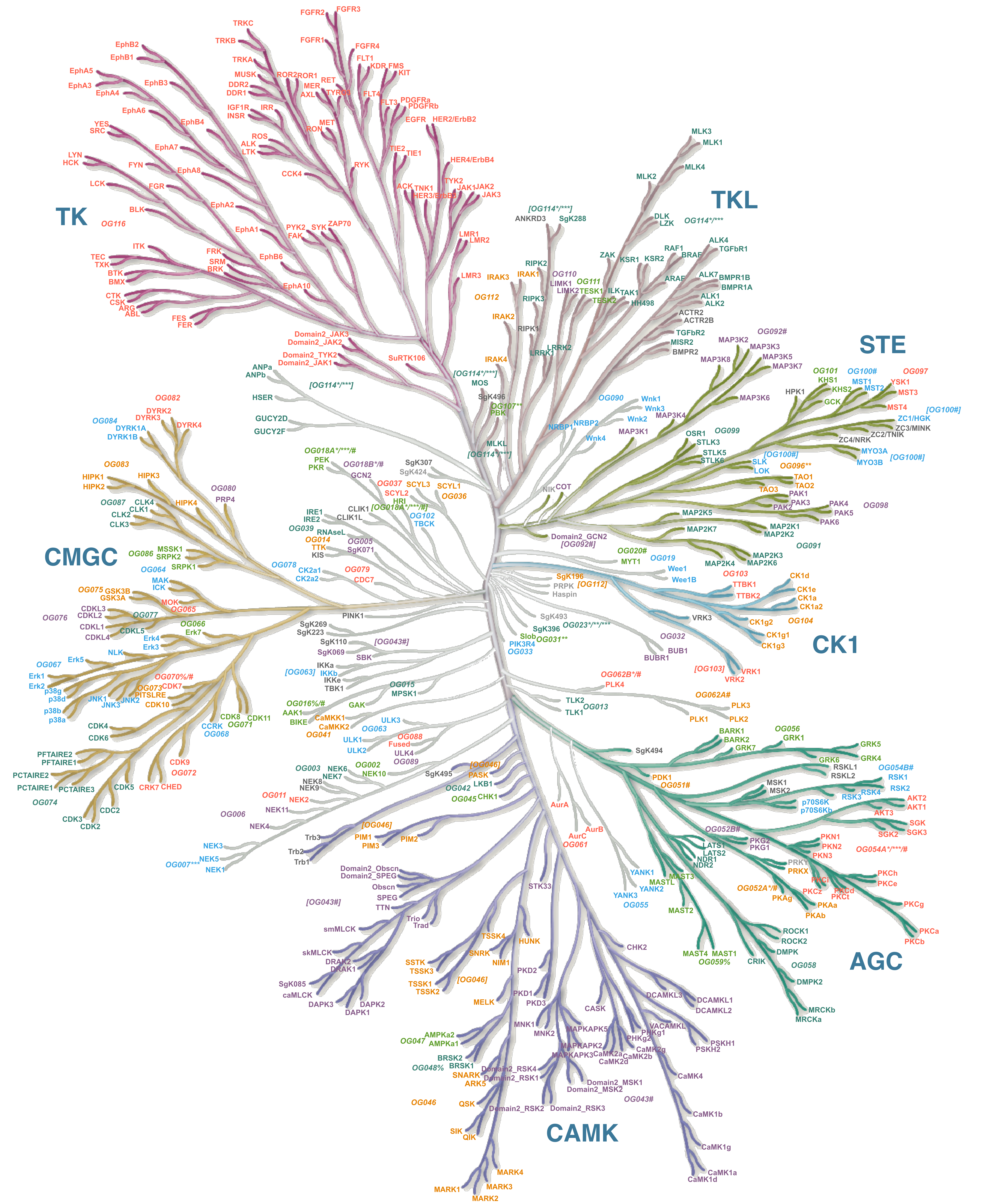


**Supplementary Fig. 2: The Human Kinome Poster[1,2], extended with LECA kinase clade assignment.** Human kinases that are assigned to the same LECA kinase clade in the eukaryotic kinome tree are indicated with the same colour and the OG number and name associated with that LECA kinase clade. LECA kinase clade OG numbers and names in-between brackets indicate LECA kinase clades that are not monophyletic in the human kinome tree. In that case, only the clade containing the human kinase with the highest bit score is indicated without brackets. Human kinases that are not assigned to any LECA kinase clade are dark grey. Human kinases that are absent in the initial ePK dataset are light grey. The meaning of special characters is the same as in Fig. 2. Illustration reproduced courtesy of Cell Signaling Technology, Inc. (www.cellsignal.com) for Kinmap[2] and adjusted with Inkscape.

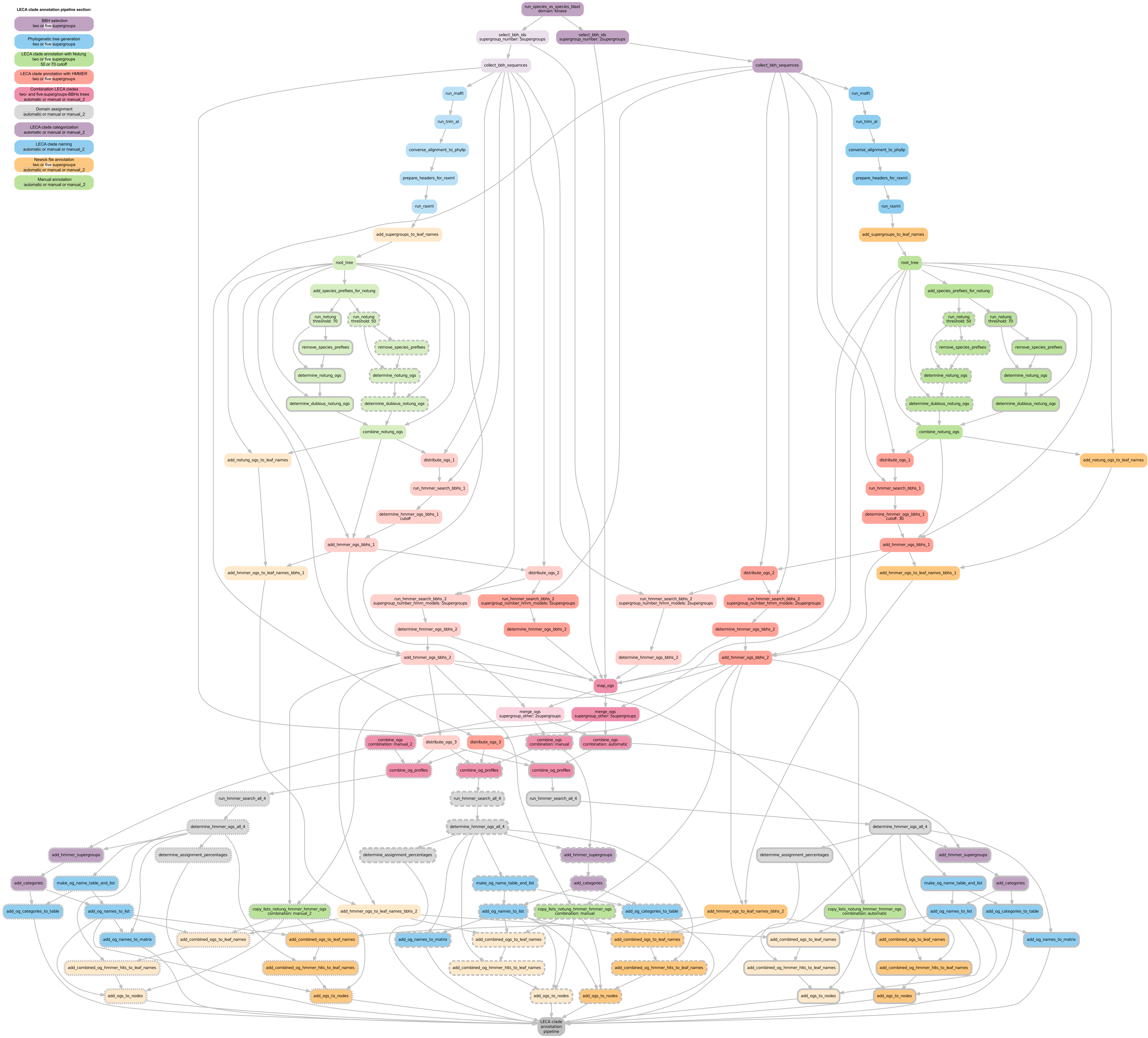


**Supplementary Fig. 3: Directed Acyclic Graph of interrelationships between input and output of rules in the LECA clade annotation pipeline Snakefile.** This figure was automatically generated with Snakemake and adjusted with Inkscape.
